# The evolution of gene regulatory programs controlling gonadal development in primates

**DOI:** 10.1101/2025.06.17.659946

**Authors:** Nils Trost, Amir Fallahshahroudi, Ioannis Sarropoulos, Céline Schneider, Julia Schmidt, Noe Mbengue, Eva Wolff, Charis Drummer, Robert Frömel, Steven Lisgo, Florent Murat, Mari Sepp, Margarida Cardoso-Moreira, Rüdiger Behr, Henrik Kaessmann

## Abstract

Sex-determining pathways produce dimorphic gonads (ovaries and testes), yet the gene regulatory programs governing gonadogenesis and their evolution in primates remain little explored. Here we report evolutionary analyses of transcriptome and chromatin accessibility data of male and female human, marmoset (New World monkey), and mouse gonadal cells spanning key prenatal stages. We find that the two primates and mouse share similar X chromosome expression dynamics, including X chromosome reactivation (XCR), and that in Klinefelter syndrome (XXY) testes, germ cells undergo female-like XCR and escape of X inactivation. New male-specific regulatory regions have emerged progressively during mammalian evolution, especially on the X following sex chromosome origination. Further analyses revealed that male-specific regulatory regions evolved faster than female-specific ones in both supporting and pre-meiotic germ cells. However, female meiotic germ cells show even higher rates of molecular evolution and exhibit a permissive chromatin state that facilitates the birth of new genes, thus resembling their adult spermatogenic counterparts. Finally, we traced both conserved and species-specific gene expression trajectories across the three mammals, uncovering candidate genes for disorders of sex development that are typically central to cell-type-specific regulatory networks. Together, our study unveils both ancestral mammalian and recently evolved gene regulatory programs that control human- and primate-specific aspects of gonadal development in both sexes.

---

Animal reproduction often relies on sex-determination mechanisms that control the development of two sexes with distinct dimorphic traits. In therian mammals (placental mammals and marsupials), expression of the Y-linked Sex-determining Region Y (SRY) gene initiates a male-specific developmental pathway^1,2^. SRY expression directs the bipotential gonads, the common gonad precursor in both sexes, to differentiate into testes in XY individuals. The absence of SRY expression in XX individuals results in ovarian development. Later, the differentiated gonads coordinate the development of sexually dimorphic traits through hormonal signalling^1,2^.

Upon sex determination, common somatic progenitor cells, which arise from the proliferation of coelomic epithelial cells ventral to the mesonephros, differentiate into Sertoli cells in males or pre-granulosa cells in females^3^, the male and female supporting cell lineages. These lineages, in turn, direct the sexual fate of the remaining gonadal somatic cells, including fetal Leydig cells in males and theca progenitor cells in females, as well as that of the primordial germ cells (PGCs) that have colonized the bipotential gonads. In males, germ cells differentiate prenatally into pre-spermatogonia and enter mitotic arrest, later forming the foundation of adult spermatogenesis. In females, germ cells differentiate into oogonia, initiate meiosis, and develop into primary oocytes, which arrest at the first meiotic prophase during fetal development. Meiosis resumes at sexual maturation and is completed only after fertilization (Fig. 1a).

**Fig. 1.**
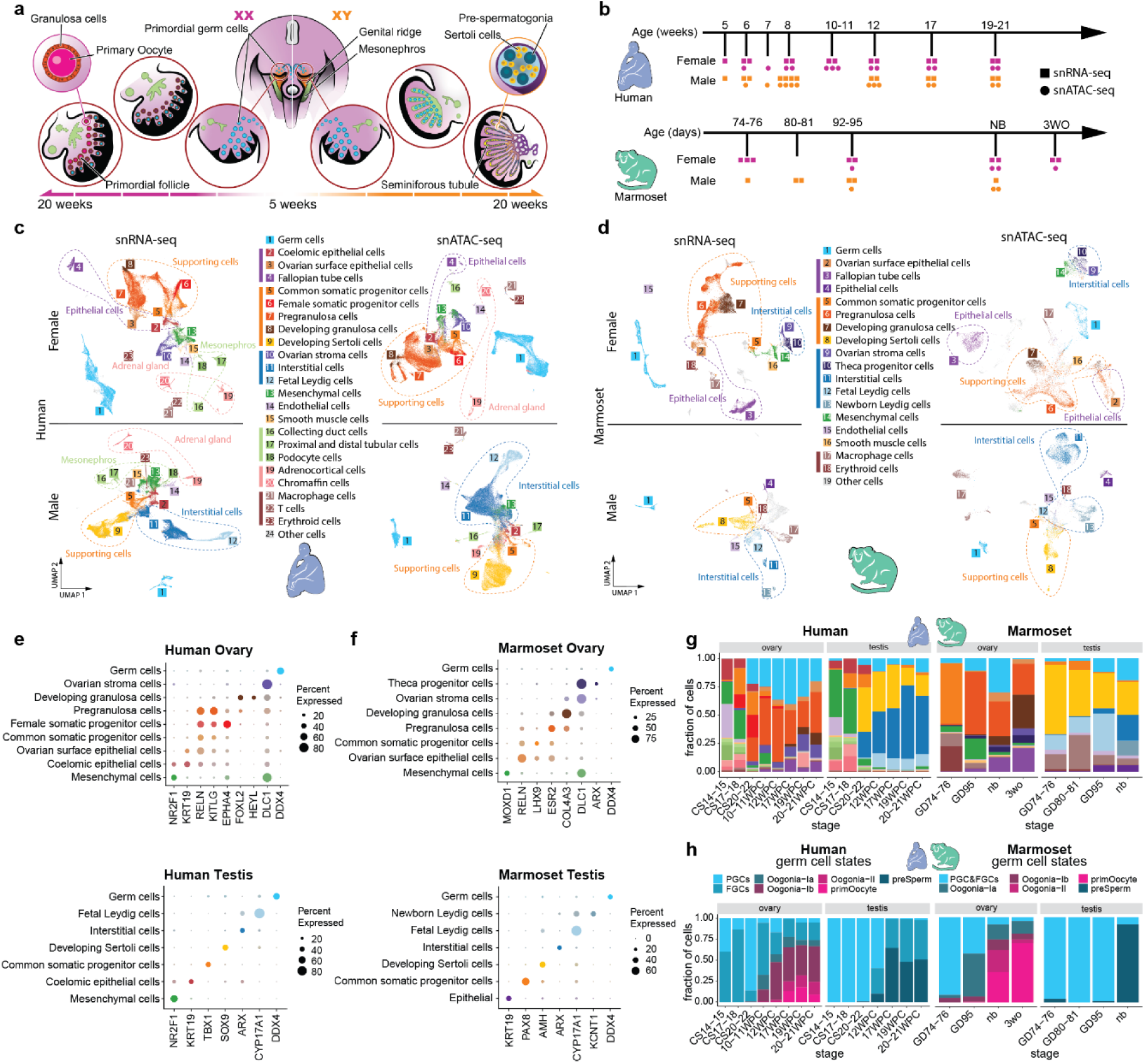
Primate atlas of gonadogenesis. **a**, Schematic of gonadogenesis and the origins of gonadal cell types. **b**, Sample collection for human and marmoset. Human-marmoset correspondences are based on cell type presence and marker gene expression. **c**, UMAPs of human ovary (top) and testis (bottom) from snRNA-seq (left) and snATAC-seq (right) datasets, labelled by cell type annotation. **d**, UMAPs of marmoset ovary (top) and testis (bottom) snRNA-seq (left) and snATAC-seq (right) datasets, labelled by cell type annotation. **e**, **f**, Marker gene expression for major cell types in the human (**e**) and marmoset (**f**) ovary (top) and testis (bottom). Dot size indicates the percentage of positive cells, colours show cell types as shown in **c** and **d**. **g**, Proportions of cell types in human and marmoset ovaries and testes across developmental stages. **h**, Proportions of male and female germ cell states in human and marmoset gonads across developmental stages.

Previous research showed that spermatogenesis in the adult testis evolves rapidly^4–10^, with meiotic spermatocytes and post-meiotic spermatids exhibiting the highest rates of molecular evolution^10–12^. However, the evolutionary dynamics of developing gonads, particularly of oogonia and oocytes, remain largely uncharacterized, leaving open the question of whether the rapid evolution of late germ cells is specific to males.

While most insights into gonadogenesis have come from mouse studies, recent single-cell sequencing studies have begun illuminating the transcriptomic^13–18^ and regulatory^17^ landscapes of human gonad development and sex determination^17^. These studies have provided insights into the emergence of gonadal cell types and their regulatory programs. However, the evolution of primate gonad development and gonadal cell types remain poorly characterized.

In this study, we generated comprehensive cell type atlases of human female and male gonad development, as well as a developmental gonad from a Klinefelter syndrome (XXY) individual, using single-nucleus RNA-seq (snRNA-seq) and single-nucleus assay for transposase accessible chromatin using sequencing (snATAC-seq). To enable evolutionary comparisons, we generated matched datasets for the marmoset, a representative of New World monkeys, which split ∼43 million years ago from the primate lineage leading to humans. Our comparative analyses of these data and published mouse datasets revealed key aspects of primate gonadal development and its evolutionary trajectory, including changes in cell type composition, gene expression dynamics, and regulatory programs. We provide an interactive online resource to explore our datasets at: https://apps.kaessmannlab.org/primate_gonadogenesis/.

## Results

### Single-cell characterisation of primate gonadal development

To define transcriptional and chromatin accessibility trajectories underlying human gonad development at single-cell resolution, we performed snRNA-seq and snATAC-seq on male and female gonadal samples spanning eight key developmental time points. These time points cover the formation and sex-fate acquisition of the bipotential gonad (5-7 weeks post conception, WPC) and its differentiation into either ovary or testis (8-21WPC). We generated 26 snRNA-seq and 26 snATAC-seq libraries, with one to four biological replicates per stage and sex (Fig. 1b, Supplementary Table 1). After filtering for high-quality cells, we retained ∼125,000 snRNA-seq profiles, with a median of 4,146 RNA molecules per nucleus, and ∼100,000 snATAC-seq profiles (median of 15,664 fragments per nucleus). To create similar data for the marmoset, we generated 13 snRNA-seq and 7 snATAC-seq libraries (Fig 1b, Supplementary Table 1) from pre- and post-natal male and female gonads (gestational day, (GD), 75 to 3 weeks old) covering gonadogenesis following sex determination, with one to three biological replicates per time point (Fig. 1b). This resulted in ∼45,000 snRNA-seq profiles (median of 2,664 RNA molecules per nucleus) and 22,000 snATAC-seq profiles (median of 21,105 fragments per nucleus) after quality filtering.

We integrated all snRNA-seq samples across time points into one gene expression atlas per sex and species using linked inference of genomic experimental relationships (LIGER)^19^. Marker gene expression within each cluster enabled us to identify all expected somatic and germ cell types and subtypes, consistent with previous cell type characterizations^17^ (Fig. 1c,d,e,f and Extended Data Figs. 1-4).

Our earliest human samples (Carnegie Stage (CS)14-18, 5-7WPC) represent the bipotential gonads and include mesenchymal cells (marked by *NR2F1* expression, *NR2F1*+), endothelial (*EGFL7*+), and coelomic epithelial cells (*UPK3B*+), common somatic progenitors (*KITLG*+), macrophages (*CD86*+), and PGCs (*POU5F1*+, *NANOG*+) (Fig. 1c,g left). We also identified cells from the neighbouring developing adrenal gland (*MC2R*+, *SOX10*+, or *DBH*+) and mesonephros (*CUBN*+, *TMEM213*+, or *NPHS2*+). After sex determination, from 8 to 21WPC, we identified both germ cells and sex-specific somatic cell types. In XX gonads, these include pre-granulosa (*FOXL2*+), ovarian stroma (*LAMA4*+), and ovarian surface epithelial cells (*KLK11*+) (Fig. 1c,g left, Supplementary Table 2). In XY gonads, we identified Sertoli (*AMH*+), interstitial (*ARX*+), and fetal Leydig cells (*CYP17A1*+) (Fig. 1c,g left, Supplementary Table 2). While all major testis cell types appeared at 8WPC, pre-granulosa cells became prevalent only after 10WPC, a result consistent with the rapid somatic cell proliferation and diversification observed during testis formation^17^.

In marmoset, we identified all post-sex determination cell types found in humans (Fig. 1d,g right, Supplementary Table 2). The gonadal cell composition of the newborn marmoset closely resembles that of a human at 20-21WPC. Additionally, in the 3-week postnatal marmoset ovaries, we identified theca cell progenitors (*GLI1*+, Extended Data Fig. 3), a cell type that emerges only after birth and is hence absent from our human dataset (Fig. 1d,f,g right).

In mouse, female germ cells are known to differentiate synchronously in an anterior-to-posterior wave, completing differentiation around embryonic day (E)16.5^20–22^. In contrast, human ovaries have a persistent gradient of differentiation even at late prenatal stages, with PGCs, fetal germ cells (FGCs), pre-meiotic oogonia (Oogonia-Ia, b), meiotic oogonia (Oogonia-II), and primary oocytes present simultaneously^17,23–26^. Our time-series snRNA-seq data confirm this asynchronous pattern in human and reveal a similar pattern in marmoset. In both species female germ cells expressing markers of PGCs (*POU5F1*+, *NANOG*+) and FGCs (*PTCHD1*+, *OPHN1*+) were detected across all sampled time points, with progressively increasing fractions of more differentiated germ cells (Fig. 1h). The asynchronous mechanism we detected in the two primates very likely represents the ancestral state because asynchronous female germ cell development is also found in outgroup species, including pigs, cattle^27,28^, and chicken^29^. This means that the synchronous wave observed in mouse likely evolved during rodent diversification.

While female germ cells enter meiosis and differentiate into primary oocytes (*NOBOX*+, *FIGLA*+), male germ cells undergo mitotic arrest and differentiate into pre-spermatogonia (*TEX41*+, *TEX15*+), which have the highest transcriptome correlation with pre-meiotic oogonia among the female germ cells (Oogonia-Ia/b, *DDX4*+, *STRA8*+) (Extended Data Fig. 5a). Our data confirm that, like female germ cells, human and marmoset male germ cells also differentiate asynchronously, with the latest sampled time points (20WPC in human, newborn in marmoset) still containing PGCs, FGCs, and pre-spermatogonia (Fig. 1h), as previously shown using transcriptome data in human^13^ and histological data in marmoset^30^. In mouse, male germ cells develop synchronously, with distinct germ cell states dominating developmental timepoints^31^. As in female germ cells, this points to a change in the regulation of germ cell development during rodent evolution.

### X chromosome dynamics and Klinefelter syndrome

Mammalian sex chromosomes emerged from ancestral autosomes^32^. This transition entailed the loss of most Y-linked genes, leaving males with a single X chromosome and prompting the evolution of dosage compensation mechanisms. Dosage compensation includes the upregulation of the X in males and in females and X chromosome inactivation (XCI, mediated by the long noncoding RNA *XIST*) in this sex^9,33,34^.

In mice, the dynamics of X chromosome expression and dosage compensation during gonadal development are well established^35^. In somatic cells, expression levels from the X chromosome are similar in males and females throughout gonadogenesis, driven by transcriptional upregulation of the single active X (Xa) in both sexes, and stable silencing of the other X (Xi) in females. Xa upregulation is also present in early germ cells from both sexes (before and during whole genome reprogramming) but is subsequently erased (i.e., absent in late germ cells). In males, this erasure leads to a permanent loss of X upregulation and, thus, of dosage compensation in late germ cells. In females, the erasure follows Xi reactivation^36–41^ (XCR) in epiblast precursors, resulting in a transient X dosage excess in early germ cells and reinstation of X dosage compensation in late female germ cells.

We found a similar dynamic of X-linked expression in human and marmoset. In human, we observed a significantly higher X-to-autosome (X:A) expression ratio in female PGCs and FGCs compared to their male counterparts (Wilcoxon rank sum test *P* = 0.000582 and *P* = 4.83×10^−5^, respectively) and compared to female somatic cells (paired Wilcoxon rank sum test *P* = 0.0007324 and *P* = 0.0002441, respectively) (Fig. 2a). Similarly, in marmoset, we saw a significantly higher X:A expression ratio in female PGC&FGCs compared to early male germ cells (Wilcoxon rank sum test *P* = 0.032), while somatic cells show no significant difference in X:A ratio between males and females (Fig. 2b). The elevated X:A ratio in early germ cell states is followed by a marked reduction in X-linked gene expression in later-stage female germ cells and pre-spermatogonia in both primates, as was also reported in mice^35^.

**Fig. 2.**
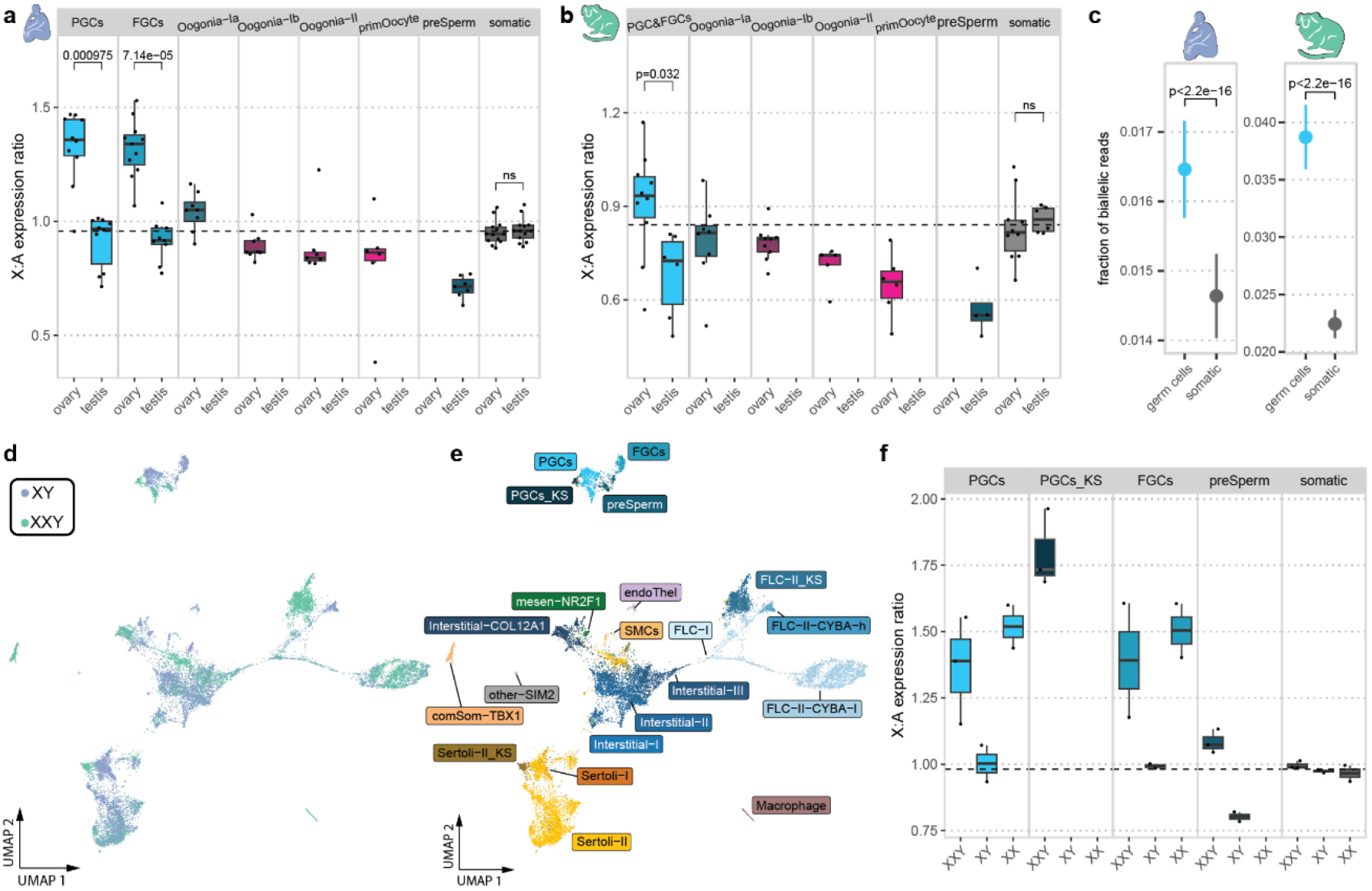
X chromosome dynamics and Klinefelter syndrome. **a**,**b**, X to autosome expression ratios in human (**a**) and marmoset (**b**) ovarian and testicular germ cell states and somatic cells. Boxes represent the first to third quartiles and whiskers extend to 1.5 times the inter-quartile range from the box hinges. Horizontal lines mark the median and dots indicate individual pseudobulk replicates. *P* values were calculated using Wilcoxon rank sum tests and adjusted for multiple hypothesis testing via the Bonferroni method. **c**, Fraction of biallelic reads in human (left) and marmoset (right) germ and somatic cells. Points indicate the mean, and ranges show the 99% confidence interval. **d**,**e**, UMAP embeddings of integrated XXY (13WPC) and XY (12WPC) snRNA-seq samples coloured by karyotype (**d**) and cell type (**e**). **f**, X to autosome expression ratios in germ cell states and somatic cells split in XXY (13 WPC), XY (12 WPC), and XX (12WPC) samples. PGC_KS denotes the Klinefelter syndrome-specific PGC cluster. In **a**, **c**, and **f**, the horizontal dashed line indicates the mean X to autosome expression ratio of all somatic cells.

Our findings differ from previous work in humans^15^. Chitiashvili and colleagues reported that female PGCs carry two active X chromosomes^15^, a finding we confirmed by measuring the fraction of biallelic expression in our human and marmoset snRNA-seq and snATAC-seq data (Wilcoxon rank sum test *P* < 2.2×10^−16^, Fig. 2c, Methods). However, their scRNA-seq data analysis showed a lower X:A ratio in PGCs compared to somatic cells, with an increase in the X:A ratio in later-stage germ cells^15^. This observation seems to contradict their finding of two active X chromosomes in PGCs, which they attribute to X chromosome dampening^15^, a transient reduction in X-linked gene expression previously described in naïve human pluripotent stem cells and pre-implantation embryos^42^.

To resolve the discrepancy between their findings and ours, we reanalyzed their dataset and confirmed the elevated X:A expression ratio in PGCs, as seen in our data and in mice^35^ (Extended Data Fig. 6a,b; see Methods for technical reasons that likely account for the reported differences). In sum, our findings do not support the occurrence of X chromosome dampening during human and marmoset germ cell differentiation and suggest that primates and mice share similar X chromosome expression dynamics during gonadal development.

Klinefelter syndrome (KS) is the most common sex chromosomal aneuploidy and is caused by an extra X chromosome in males (47, XXY)^43^ and is associated with highly variable symptoms. Aberrant X chromosome dosage compensation has been proposed to contribute to partial or complete loss of fertility^44^. XCI escapees also contribute to the excess of X-linked gene expression in XXY individuals^43,45^ and this escape has been associated with KS and comorbidities in adult patient data^44^. However, despite previous investigations in KS mouse models^35,46^ and humans^47^, they remain uncharacterized in humans at a single-cell level.

We investigated a human 13WPC KS testis using snRNA-seq (8,963 nuclei) and snATAC-seq (11,022 nuclei) across three technical replicates. Data annotation and integration with 12WPC testis samples revealed all expected cell types for this developmental stage. XXY cells integrated well, except for three XXY-specific clusters expressing markers of fetal Leydig cells (annotated as FLC-II_KS), Sertoli cells (Sertoli-II_KS), and germ cells (PGCs_KS), respectively (Fig. 2d,e and Extended Data Fig. 6c,d). The XXY-specific fetal Leydig cell cluster exhibited elevated expression of *CYBA*, a gene typically expressed in later stages (17–19WPC). Consistent with studies in adult tissues^48^, somatic XXY cells expressed *XIST* (Extended Data Fig. 6e). Analyses of differentially expressed genes across karyotypes revealed an enrichment of known XCI escapees (i.e., *RPS4X*, *XIST*, *IL3RA*, *JPX* and *ASMTL*) among genes upregulated in XXY compared to XY somatic cells (n = 5, *P* = 0.003413, Fisher’s exact test; Extended Data Fig. 6f). *RPS4X* and *JPX,* two of the only three consistent escapees located on the long arm^45^, along with *XIST,* are significantly more highly expressed in the somatic cells of ovaries than in testes at the same developmental stage (12WPC), suggesting that the somatic cells of XXY testes exhibit a female-like XCI escape profile (Extended Data Fig. 6g), in line with observations in the kidney and liver of adult mouse models of KS^49^.

Somatic XXY cells showed X:A ratios comparable to those in XY testes and XX ovary somatic cells, indicating functional XCI in the developing KS testis (Fig. 2f), consistent with studies in adult tissues^50–52^. In contrast, XXY PGCs and FGCs exhibited elevated X:A ratios for protein-coding genes (Fig. 2f), along with an increased number of chromatin accessibility peaks (snATAC-seq) on the X chromosome (Extended Data Fig. 6h), mirroring patterns observed in XX germ cells. These features suggest there is X chromosome reactivation (XCR) in KS PGCs/FGCs, as seen in mouse models^35,46^. Notably, unlike XY pre-spermatogonia, which show a reduced X:A ratio compared to their PGC/FGC precursors and somatic cells (see above), XXY pre-spermatogonia maintain X dosage (Fig. 2f). Collectively, these observations support the notion that X chromosome number, not phenotypic sex, determines X chromosome dynamics during germ cell differentiation^35,53^.

### Changing chromatin landscape during sex differentiation

Sex chromosome differences between males and females trigger a cascade of transcription factors that drive the developing gonad towards either male or female fate^1,2,54–57^. To better understand this process, we assessed chromatin changes of differentiating supporting and germ cells between sexes. We transferred labels from our annotated snRNA-seq data to the corresponding snATAC-seq datasets based on gene scores calculated from chromatin accessibility profiles and identified peaks of accessibility as a proxy for cis-regulatory elements (CREs) using ArchR^58^ (Extended Data Fig. 7a). For female-to-male comparisons, we aggregated the human snATAC-seq data by developmental time points (CS 17-22, 10-12WPC, 17-19WPC, and 20-21WPC) for supporting cell types and by three differentiation states (PGCs, FGCs, and pre-meiotic Oogonia-Ia or pre-spermatogonia) that can be compared between the sexes for germ cells. We classified CREs in these groups based as male-specific, female-specific, or shared. We excluded the sex chromosomes for the sex-based comparisons for better comparability between the sexes (see Methods, Extended Data Fig. 7b,c).

In mice^59^, supporting cells show a trend from shared CREs to sex-specific CREs during gonadal development. Our human data mirror this trend (Fig. 3a, left, Extended Data Fig. 7b) but also reveal a shift from more sex-specific CREs in female supporting cells in early stages to more sex-specific CREs in male supporting cells in later stages (Fig. 3a, left). Shortly after sex differentiation (CS17-22), supporting cells show a high proportion of female-specific CREs (Fig. 3a, left), a surprising result given that supporting cell differentiation occurs earlier in males^60^. Similarly, female PGCs exhibit many sex-specific CREs, while FGCs display more shared CREs (Fig. 3a, right).

**Fig. 3.**
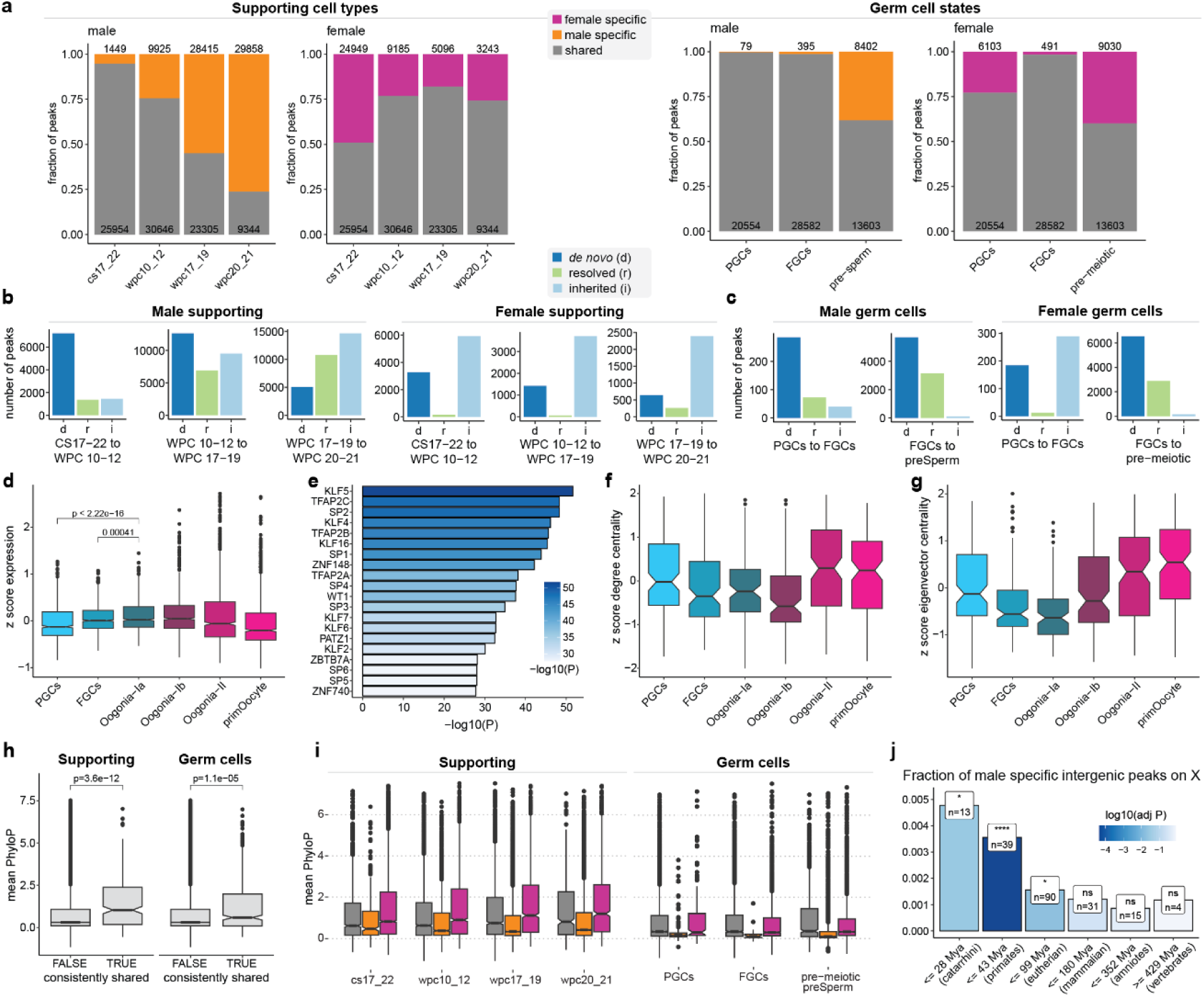
Chromatin landscape during human gonadogenesis. **a**, Fractions of shared and sex-specific autosomal peaks per stage in the supporting lineage (left) and per cell state in the germ cell lineage (right) of the human developing gonad. Sub panels show the fractions for peaks for male and female cells. Absolute numbers of peaks are shown above (sex-specific) and below (shared). **b**,**c**, CRE dynamics in the human female and male supporting cells (**b**) and germ cells (**c**). Sub panels show the numbers of *de novo*, resolved, and inherited peaks from one stage to the next. **d**, Relative expression (z-score) of putative target genes of de-novo acquired peaks in pre-meiotic oogonia per germ cell state, based on pseudobulk replicates. *P* values were calculated using a Wilcoxon rank sum test. **e**, Top 20 TFs assigned to enriched motifs in de novo acquired peaks in pre-meiotic oogonia. The length and colour of the bars represent the negative decadic logarithm of the adjusted (Benjamini-Hochberg) *P* value of Fisher’s exact test measuring the enrichment of the motif in the peaks. **f**, **g**, Relative degree centrality (**f**) and the relative eigenvector centrality (**g**) of TFs assigned to enriched motifs in de-novo acquired peaks in pre-meiotic oogonia in the GRN of each germ cell state. **h**, Mean phyloP scores of dynamic or consistently shared peaks in the supporting and the germ cell lineage. *P* values were calculated using a Wilcoxon rank sum test. **i**, Mean phyloP scores of dynamic shared and sex-specific peaks per stage in the supporting and the germ cell lineage. **j**, Fraction of male-specific intergenic peaks in supporting cells on the X chromosome in each age group. Bar colours show the negative decadic logarithm of the adjusted (Benjamini-Hochberg) *P* values (Fisher’s exact test) of the overrepresentation of male-specific peaks on the X chromosome. Mya, million years ago.

In supporting cells, most male-specific CREs are acquired de novo (region inaccessible in both sexes in the previous stage(s)) from CS17-22 to 10-12WPC, with fewer inherited or resolved CREs (i.e., previously shared CREs becoming sex-specific) (Fig. 3b, left). This trend continues in later stages, with increasing proportions of resolved and especially inherited male-specific CREs. These results indicate that the male-specific chromatin landscape is predominantly shaped by the opening of new regions and retention of shared accessible regions that close during female development. In contrast, in female supporting cells most sex-specific CREs are inherited throughout development, with many already established by CS17-22. There are fewer de novo CREs and almost no resolved CREs (Fig. 3b, right). These findings suggest that the female chromatin landscape is established early in development and remains stable thereafter.

Male germ cells show a pattern of predominantly de novo male-specific CREs from PGCs to FGCs and from FGCs to pre-spermatogonia (Fig. 3c, left). In contrast, the female PGCs to FGCs transition is marked by inherited female-specific CREs, with a considerable increase in de novo female-specific CREs in pre-meiotic oogonia (Fig. 3c, right). To explore the function of these *de novo* regions, we built cell state and sex-specific gene regulatory networks (GRNs) and linked peaks to their putative target genes (Methods). We identified 1128 putative target genes of de novo female-specific regions in pre-meiotic oogonia and 1002 for male-specific regions in pre-spermatogonia, with significant Gene Ontology (GO)^61,62^ term enrichment related to the cell cycle and meiosis. Target genes of *de novo* female-specific regions, including *STRA8*, showed increased expression leading up to meiosis, peaking in Oogonia-Ib (Fig. 3d). The transcription factors (TFs) matching the ∼400 enriched TF motifs in the de novo female-specific CREs of premeiotic oogonia showed the highest connectivity in the GRNs of meiotic oogonia and primary oocytes (Fig. 3e-g), suggesting that these regions represent early chromatin changes associated with the start of meiosis.

Previous work has highlighted considerable enhancer pleiotropy, especially during early development^63,64^. In our data, we found intergenic CREs that remain consistently shared between sexes. These shared CREs are often found in both the supporting and germ cell lineages (*P* < 2.2×10^−16^, Fisher’s exact test) and are linked to 196 target genes enriched in broad GO terms like multicellular organism development. Notably, *GATA4*, *CHD7*, and *PSMC3IP*, genes associated with disorders of sex development (DSDs)^65–67^, are among the targets. Evolutionary analysis showed that shared CREs are under higher sequence constraint than dynamic and sex-specific CREs (Wilcoxon rank sum test *P* = 0.0011) (Fig. 3h), indicating their regulatory importance across the sexes. Male-specific CREs have lower sequence constraints than female-specific and shared CREs in supporting cells as well as the pre-meiotic germ cells compared here (Fig. 3i), pointing to an overall faster evolution of male gene regulatory programs.

Further analyses suggest that new male-specific CREs have continuously arisen in pre-spermatogonia and especially in later supporting cells during mammalian evolution (Extended Data Fig. 7d,e), in particular on the X chromosome since the origination of therian sex chromosomes (Fig. 3j, Extended Data Fig. 7f). The latter finding aligns with the accumulation of genes expressed in male cell types on the X chromosome^6,10,68,69^.

### Developmental gene expression trajectories

We next sought to investigate the dynamics of gene expression during gonad development, infer the profiles of genes associated with disorders and compare these patterns across species. We created pseudotime trajectories for human female and male germ cells and the four main somatic lineages using Monocle 3^70–72^ (Extended Data Fig. 8a-g). We identified genes with dynamic expression using a negative binomial generalized additive model and clustered them based on their expression trajectories (Extended Data Fig. 8c). This analysis highlighted key transitions in germ cells, for instance clusters 1, 6, and 8 contains genes expressed in PGCs and FGCs (e.g., *POU5F1*), and cluster 9 includes genes expressed in meiotic oogonia (e.g., *SPO11*). In somatic cells, early developmental clusters 1 and 5 showed expression of genes (e.g., *KRT19* and *UPK3B)* enriched in mitotic cell cycle functions (Extended Data Fig. 8g).

We associated 35 known DSD genes with specific male or female somatic or germ cell differentiation trajectories (adjusted *P* value < 0.05, Wald test; Extended Data Fig. 8d,h and Supplementary Table 3). For example, *ZFPM2*, linked to gonadal dysgenesis, was expressed in pre-spermatogonia (Extended Data Fig. 8d) and previously reported only in mouse developing Sertoli cells^73^. Notably, its co-factor GATA4 was recently shown to have primate-specific expression in germ cells^17^. Another example is *MAMLD1*, which is associated with hypospadias^72^. We observed *MAMLD1* upregulation in Sertoli cells and fetal Leydig cells (Extended Data Fig. 8h), in line with previous findings in mouse and human^74,75^. Additionally, we observed expression in pre-spermatogonia, which was not previously reported (Extended Data Fig. 8d).

Given that much of the understanding of gonadogenesis is based on mouse models, we compared our human and marmoset data with a published mouse scRNA-seq dataset for germ cells from embryonic day (E) 12.5 through E16.5^76^ and somatic cells from E10.5 through E16.5^77^.

To study female germ cell differentiation, we integrated the datasets from human, marmoset, and mouse (13,064, 4,586, and 7,554 germ cells, respectively) based on 1:1 orthologous genes and created pseudobulks along the pseudotime trajectory of germ cell differentiation (Fig. 4a). We verified alignments of pseudotime trajectories using marker genes known for their roles in germ cell development, (e.g., *STRA8*, *NOBOX*, and *ZP3*). We confirmed conserved expression patterns previously identified^17^, including those of *DMRTC2*, *ZNF711*, and *DMRTB1* after meiosis initiation, *PBX3* in Oogonia-II, and *TP63* and *ZHX3* in primary oocytes (Extended Data Fig 8i).

**Fig. 4.**
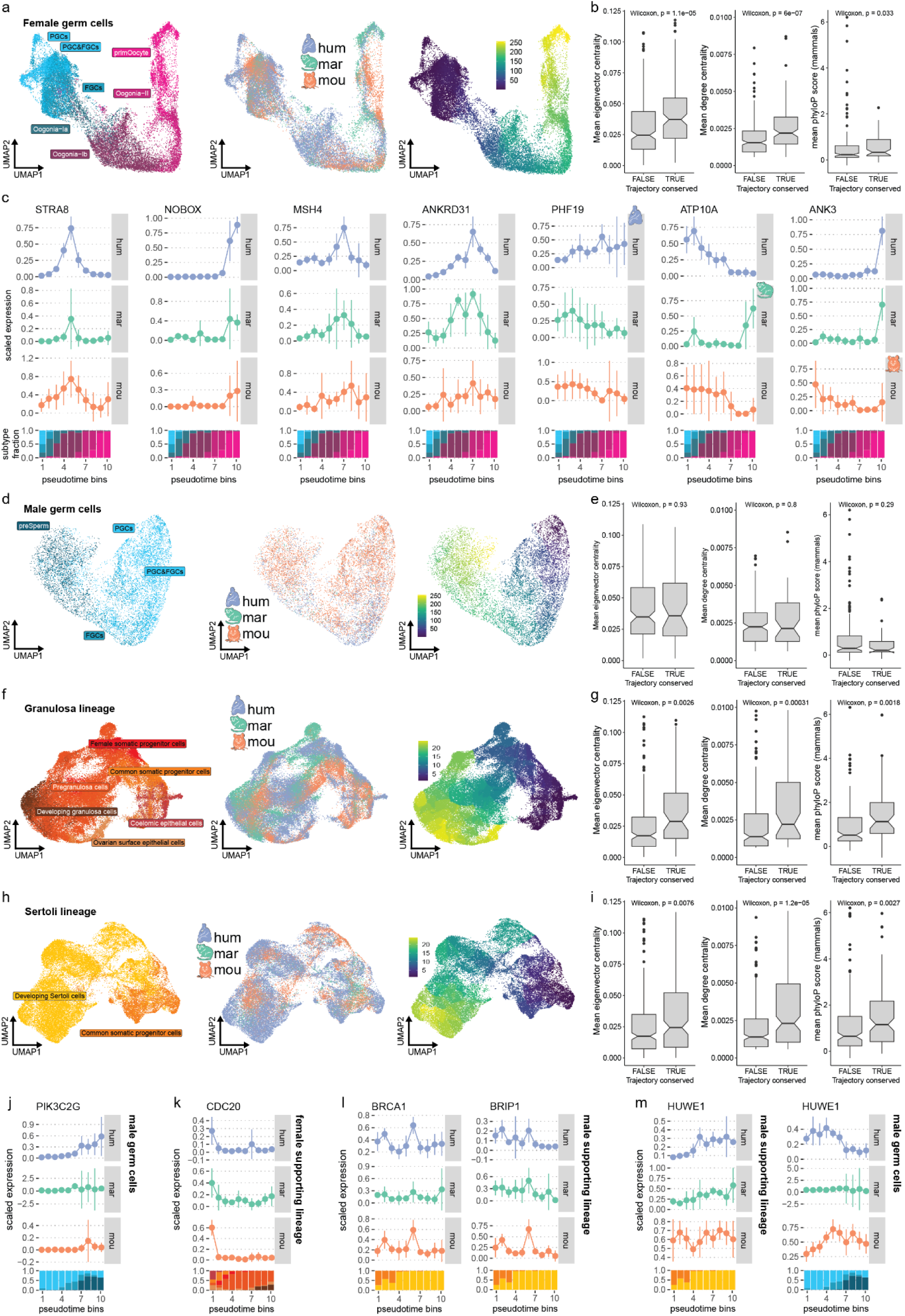
Developmental gene expression trajectories. **a**, Joint snRNA-seq UMAP embeddings of female human, marmoset, and mouse germ cells coloured by germ cell state (left), species (centre) and pseudotime value (right). **b** Connectedness of genes with conserved or changed trajectories between human, marmoset, and mouse female germ cells (left: eigenvector centrality, centre: degree centrality) and sequence constraint (phyloP scores) of distal CREs associated to genes with conserved or changed trajectories (right). **c,** Examples of genes with shared gene expression trajectories (*STRA8*, *NOBOX*, *MSH4*, *ANKRD3*), human specific trajectories (*PHF19*), marmoset specific trajectories (*ATP10A*) and mouse specific trajectories (*ANK3*) in female germ cells. Point range plots show mean scaled expression and 95% confidence intervals in each pseudotime bin. Filled bar plots show the contribution of germ cell states to each pseudotime bin. **d,** Joint snRNA-seq UMAP embeddings of male human, marmoset, and mouse germ cells coloured by germ cell state (left), species (centre) and pseudotime value (right). **e,** Connectedness of genes with conserved or changed trajectories between human, marmoset, and mouse male germ cells (left: eigenvector centrality, centre: degree centrality) and sequence constraint (phyloP scores) of distal CREs associated to genes with conserved or changed trajectories (right). **f,** Joint snRNA-seq UMAP embeddings of female human, marmoset, and mouse supporting cells coloured by cell type (left), species (centre) and pseudotime value (right). **g,** Connectedness of genes with conserved or changed trajectories between human, marmoset, and mouse female supporting cells (left: eigenvector centrality, centre: degree centrality) and sequence constraint (phyloP scores) of distal CREs associated with genes with conserved or changed trajectories (right). **h,** Joint snRNA-seq UMAP embeddings of male human, marmoset, and mouse supporting cells coloured by cell type (left), species (centre) and pseudotime value (right). **i,** Connectedness of genes with conserved or changed trajectories between human, marmoset, and mouse male supporting cells (left: eigenvector centrality, centre: degree centrality) and sequence constraint (phyloP scores) of distal CREs associated to genes with conserved or changed trajectories (right). **j, k, l, m,** Trajectory comparisons of selected genes in supporting and germ cell lineages accross human, marmoset and mouse.

We identified 486 genes with conserved dynamic temporal trajectories across the three species in female germ cells (Fig. 4c, Supplementary Table 4). These include well-known markers of germ cell differentiation, such as *STRA8*, *SPO11*, *FIGLA*, and *NOBOX*. Genes associated with human DSDs, abnormal fertility in mice, and sex disorder-related variants (see Methods) were significantly enriched among these genes (*P* = 4.4×10^−6^, *P* = 0.0019, and *P* < 2.2×10^−16^, respectively, Fisher’s exact test), highlighting their important roles in germ cell differentiation. Examples include *MSH4* and *ANKRD31*, which are linked to premature ovarian failure or insufficiency, with expression peaking in Oogonia-II in all studied species. We also found that genes with conserved trajectories are more connected in germ cell GRNs (Wilcoxon rank sum test *P* = 1.1×10^−5^ and *P* = 6×10^−7^, Fig. 4b) than genes with divergent trajectories, possibly reflecting regulatory roles. Genes with conserved trajectories are also significantly more frequently associated with conserved distal CREs (Wilcoxon rank sum test *P* = 0.033, Fig. 4b).

In addition to genes with conserved expression during female germ cell development, we identified 688 genes with primate- or mouse-specific expression trajectories (e.g., *ANK3*), 276 with human-specific expression trajectories (e.g., *PHF19*), and 535 with marmoset-specific expression trajectories (e.g., *ATP10A*) (Fig. 4a, Supplementary Table 4).

We next conducted similar analyses for male germ cells from human, marmoset, and mouse (3,297, 458, and 6,502 germ cells, respectively), covering differentiation from PGCs to pre-spermatogonia (Fig. 4d). We verified the alignment of the pseudotime trajectories using marker genes (e.g. *TUBB4B*, *DDX4,* and *TEX15;* Extended Data Fig. 8j). We confirmed *SOX4* expression in PGCs and pre-spermatogonia of humans and mice^17^, and also found it in marmoset PGCs and pre-spermatogonia (Extended Data Fig. 8j). Across the three species, we identified 211 genes with conserved trajectories and genes that evolved species-specific trajectories (human-specific: 202, marmoset-specific: 368, primate- or mouse-specific: 358; Supplementary Table 5) in male germ cells. Genes with conserved trajectories were significantly enriched among genes with pathogenic variants linked to sex-related disorders (*P* = 2.2×10^−16^, Fisher’s exact test), including *PIK3C2G*, expressed in pre-spermatogonia (Fig. 4j), which is annotated with nine variants related to spermatogenic failure. DSD genes are not significantly enriched among genes with conserved trajectories (*P* = 0.064, Fisher’s exact test). We found no significant differences in GRN connectivity or CRE sequence conservation between genes with conserved and divergent trajectories in male germ cell differentiation (Fig. 4e).

We also examined the evolution of gene expression trajectories in granulosa cell differentiation (human: 25,286 cells, marmoset: 15,654 cells, and mouse: 14,843 cells) and Sertoli cell differentiation (human: 23,171 cells, marmoset: 2,033 cells, and mouse: 8,045 cells) in the three species (Fig. 4f,h). Early coelomic epithelial cells were excluded from the male analyses due to insufficient data for the marmoset. We confirmed previous findings^17^, including the downregulation of *UPK3B* and *LRRN4* and upregulation of *WNT6* in early common somatic progenitor cells across all species (Extended Data Fig. 8k,l). Additionally, we observed *UPK3B* and *LRRN4* upregulation in later granulosa states, which was not previously reported. We found species differences in *WNT6* expression, which peaks in early Sertoli and pregranulosa cells before a temporary downregulation in human and marmoset, but not in mice (Extended Data Fig. 8k,l). In addition to *TSPAN8* expression in human early common somatic progenitor cells^17^, we detected its expression in mouse coelomic epithelial cells (with downregulation followed by upregulation in pregranulosa and developing granulosa cells) and observed a gradual expression increase during granulosa cell development in marmoset (Extended Data Fig. 8l).

We identified 348 genes in the female and 623 genes in the male somatic lineages with conserved trajectories alongside species-specific changes (human: 258 and 366, marmoset: 546 and 477, primate/mouse: 461 and 507) (Supplementary Tables 6 and 7). Genes with conserved trajectories are more connected in their GRNs (eigenvector centrality and degree centrality; granulosa: *P* = 0.0026 and *P* = 0.00031, Sertoli: *P* = 0.0076 and *P* = 1.2 × 10^−5^, Wilcoxon rank sum test; Fig. 4g,i) and linked to more conserved distal CREs (*P* = 0.0018 and *P* = 0.0027 in granulosa and Sertoli lineages, respectively, Wilcoxon rank sum test; Fig. 4g,i). We found a significant enrichment of genes with pathogenic sex-related variants among conserved granulosa cell trajectories (Fisher’s exact test *P* = 2.848 × 10^−5^). These include *CDC20*, expressed in coelomic epithelial cells and ovarian surface epithelial cells and linked to oocyte maturation defects and infertility (Fig. 4k). In Sertoli cells, genes with conserved trajectories are enriched among DSD genes, fertility-related genes, and sex-related pathogenic variants (Fisher’s exact test *P* = 0.00096, *P* = 0.018, and *P* = 9.09 × 10^−6^, respectively). Notably, *BRCA1* and *BRIP1*, associated with ovarian abnormality and cancer but – so far – not testis disorders, showed conserved expression in early Sertoli cells (Fig. 4l).

Among the genes with either primate- or mouse-specific trajectories in Sertoli cells, we identified *HUWE1* (Fig. 4m), an X-linked E3 ubiquitin ligase associated with male infertility, including azoospermia and oligozoospermia^78,79^. We also found that this gene’s expression trajectory in prenatal germ cells is reversed between humans and mice (Fig. 4m), suggesting differences in gene regulation between primates and mice, and therefore limitations in using mouse models to study this gene’s role in human development.

### Gonadal cell type evolution

We broadened our analysis from individual genes to cell types in developing gonads to understand their evolutionary dynamics. To do so, we assessed the levels of sequence constraint in accessible intergenic CREs using phastCons^80^ and phyloP^68^ scores and in protein-coding sequences using the normalized rate of amino acid-altering substitutions (dN/dS). We also estimated the evolutionary age of the transcriptome of each cell type using the phylogenetic age of the expressed genes^5,81,82^ and the fractions of recently duplicated genes.

In somatic cell types, CRE and coding sequence constraints decrease progressively during development while the expression of young genes increases, a pattern common to other organs^5,8,64^ (Fig. 5a,d,e). We observed an increase in tissue- and time-specific gene expression as development progresses, suggesting progressively lower expression pleiotropic constraints^5^ (Fig. 5b). Together, these observations indicate a reduction in purifying selection (functional constraint) over development, facilitating an increase in evolutionary innovations, such as the emergence of new genes^5^.

**Fig. 5.**
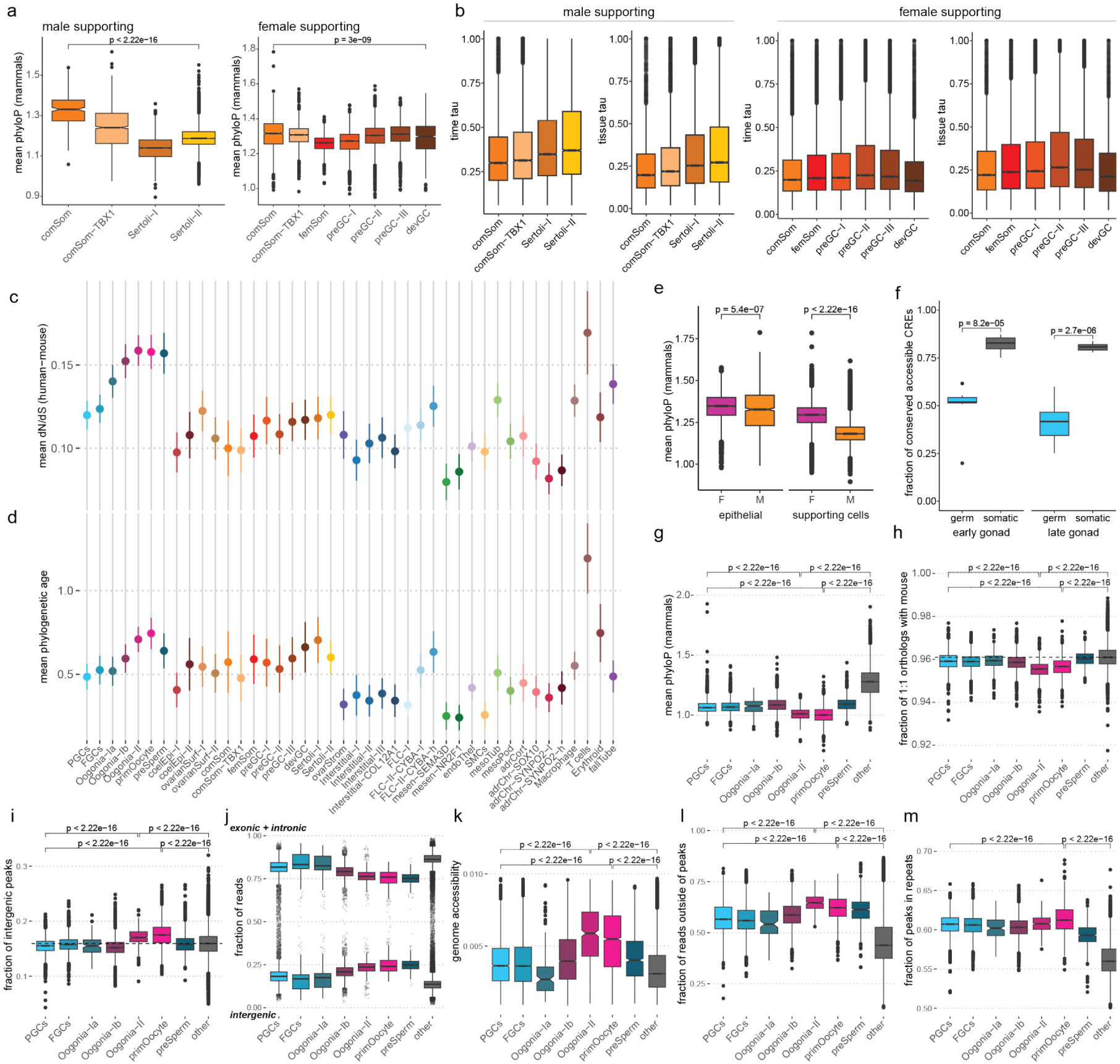
Gonadal cell type evolution. **a**, Sequence constraint of distal CREs in human male (left) and female supporting lineages (right). **b,** Time and tissue specificity of expressed genes in the human granulosa cell lineage subtypes. **c,** dN/dS values between human and mouse of human marker genes per subtype. Points show the mean; ranges show 95% confidence intervals. **d,** Phylogenetic age of human marker genes per subtype (higher values correspond to younger genes). Points show the mean; ranges show the 95% confidence intervals. **e,** Sequence constraint of accessible distal CREs of human female and male coelomic epithelial and supporting cells (granulosa and Sertoli cells, respectively). *P* values calculated by Wilcoxon rank sum tests. **f,** Fraction of CRE with conserved activity between human and marmoset in early germ and somatic cells (human: 10-12WPC, marmoset: GD92-95) and late germ and somatic cells (human: 19-21WPC, marmoset: newborn). *P* values calculated by Wilcoxon rank sum tests. **g**, Sequence constraint of accessible distal CREs of human germ cell states and all other gonadal cells. Benjamini-Hochberg adjusted *P* values calculated by Wilcoxon rank sum tests. **h**, Fraction of 1:1 orthologs between human and mouse among expressed genes in human germ cell states and all other gonadal cells. The dashed horizontal line shows the median of the somatic cell types. Benjamini-Hochberg adjusted *P* values calculated by Wilcoxon rank sum tests. **i,** Fraction of intergenic snATAC-seq peaks in human germ cell states and all other gonadal cells. The dashed horizontal line shows the median of the somatic cell types. Benjamini-Hochberg adjusted *P* values calculated by Wilcoxon rank sum tests. **j,** Fraction of genic (top) and intergenic (bottom) snRNA-seq reads in human germ cell states and all other gonadal cells. **k**, Global genome accessibility in human germ cell states and all other gonadal cells. Benjamini-Hochberg adjusted *P* values calculated by Wilcoxon rank sum tests. **l**, Fraction of snATAC-seq reads outside of peaks in human germ cell states and all other gonadal cells. Benjamini-Hochberg adjusted *P* values calculated by Wilcoxon rank sum tests. **m**, Fraction of snATAC-seq peaks overlapping repetitive elements in human germ cell states and all other gonadal cells. Benjamini-Hochberg adjusted *P* values calculated by Wilcoxon rank sum tests.

The slowest evolving gonadal cells are chromaffin cells, a neuroendocrine cell type, with the highest coding sequence constraint and the evolutionarily oldest transcriptome (Fig. 5c,d and Extended Data Fig. 9a,b). Comparisons with developmental snATAC-seq data for the human cerebral cortex^83^ show that the sequence conservation of CREs in chromaffin cells is similar to that of neuronal cell types of the developing brain, a slowly evolving organ at the molecular level^4–6^ (Extended Data Fig. 9a). In contrast, the fastest-evolving cells are immune cells, which are known to display high rates of molecular evolution^64,84–86^, and germ cells (Fig. 5c,d).

Intergenic CREs in male supporting cells (developing and fetal Sertoli cells) exhibit lower constraints than their female counterparts (granulosa cells) (Fig. 5e). Germ cells from both sexes overall show significantly lower conservation of CRE accessibility between human and marmoset than somatic cells, an effect that increases in later stages of development (Fig. 5f). Unexpectedly, contrary to earlier (i.e., pre-meiotic) germ cells (see above; Fig. 3i), meiotic oogonia and primary oocytes show significantly lower CRE sequence conservation than all other germ cells and supporting cell types, and they also exhibit less coding sequence constraint and younger transcriptomes (Fig. 5c,d,g and Extended Data Fig. 9b). Moreover, meiotic oogonia and primary oocytes exhibit the lowest fraction of expressed 1:1 orthologs between human and mouse, suggesting a high fraction of recently duplicated genes in these cell types (Fig. 5h). Notably, prior studies indicated that male meiotic germ cells (spermatocytes) and especially post-meiotic germ cells (round spermatids) during adult spermatogenesis evolve rapidly^10,11^. While the female equivalents of round spermatids (i.e., post-fertilization oocytes, ootids) are virtually inaccessible for analyses in humans, our observation that coding sequence constraints and transcriptome ages in primary oocytes are as low as those of zygotene spermatocytes in adult human testes is remarkable (Extended Data Fig. 9c,d). The developing marmoset and mouse ovary show a consistent pattern, with genes expressed in meiotic oogonia having significantly lower coding-sequence constraints than those expressed in earlier germ cell states and somatic cells (Extended Data Fig. 9e,f). These findings suggest that the rapid evolution of meiotic germ cells is a shared feature of both male and female gametogenesis (and not male-specific as previously believed^4–10^) and that this pattern is conserved across mammals.

The enrichment of recently evolved (new) genes in meiotic and post-meiotic spermatogenic cells was attributed to a transcriptionally permissive chromatin landscape that facilitates new gene origination^10,11,87^. To investigate whether this mechanism also explains the young transcriptomes of meiotic oogonia (Oogonia-II) and primary oocytes, we assessed chromatin accessibility and the extent of transcription across the genome in all human pre-natal germ cells and somatic cells by measuring the fraction of intergenic snATAC-seq peaks and intergenic snRNA-seq reads. This analysis revealed an increase in intergenic chromatin accessibility and transcription during germ cell development (Fig. 5i,j). Consistently, female germ cells in the marmoset gonad show an increase in intergenic transcription throughout development with a peak in primary oocytes (Extended Data Fig. 9g). The intergenic snRNA-seq reads in human primary oocytes and pre-spermatogonia are, on average, located further away from genes (both up- and downstream) than in other cells (Extended Data Fig. 9h), suggesting the high fraction of intergenic reads in these cell types is not due to transcriptional readthrough. Moreover, we found that human Oogonia-II and primary oocytes have the highest fractions of open chromatin (Fig. 5k), snATAC-seq reads outside of peaks (Fig. 5l), and snATAC-seq reads within repeats (Fig. 5m), suggesting noisy accessibility patterns across the genome in these cell types. Together, our analyses suggest a permissive chromatin state in late (meiotic) prenatal female germ cells that likely facilitated the emergence of new genes akin to what has been described for late (meiotic) germ cells in adult males^10,11,87^.

## Discussion

Our study provides comprehensive snRNA-seq and snATAC-seq datasets covering male and female pre-natal gonadogenesis in human and marmoset. To facilitate exploration of our data, we developed an online resource: https://apps.kaessmannlab.org/primate_gonadogenesis/.

Our analyses show that asynchronous germ cell development is likely a shared feature of primate gonad development. In combination with previous findings in other species, our results suggest that the synchronous germ cell differentiation mode observed in mice is a derived feature. Future studies including a broader range of species could help clarify the evolutionary origin of this developmental mode.

Our germ cell data showed that XCR occurs in human female PGCs and FGCs, leading to a transiently increased X:A ratio during germ cell differentiation. Using the first snRNA-seq and snATAC-seq datasets of a prenatal XXY KS testis sample, we showed that XCI and XCR occur in human XXY germ cells and identified female-like XCI escapee expression patterns. These results advance our understanding of the molecular basis of fertility issues in KS.

By comparing chromatin changes throughout development and between male and female gonads, we discovered that the female-specific chromatin landscape is established early after sex determination, while male-specific regulatory regions emerge progressively throughout development. This occurs either de novo or through the resolution of previously shared CREs in males. New male-specific accessible regions have formed continuously during mammalian evolution and have evolved faster than female-specific or sex-shared CREs in supporting cells and pre-meiotic germ cells, suggesting rapid gene regulatory evolution in these male cell types. In particular, the X chromosome has accumulated male-specific CREs throughout evolution, likely reflecting the preferential emergence of genes specifically expressed in male cells on this chromosome during evolution^6,10,68,69^.

We characterized gene expression trajectories during human gonadogenesis of the major somatic and germ cell lineages, linked DSD genes to specific lineages, and described their temporal expression. Genes with conserved expression trajectories between mice and primates are associated with denser GRNs and more conserved regulatory elements. They are also enriched for known DSD genes, pathogenic variants associated with gonadal disorders and genes linked to abnormal fertility. These results suggest that the genes with conserved trajectories are crucial in gonadogenesis, and highlight genes not yet linked to DSDs as promising candidates for future studies. Additionally, genes with divergent trajectories between mice and primates provide a resource for evaluating the transferability of insights from mouse models.

Finally, we showed that the pattern of rapid evolution of male meiotic cell types extends to female meiotic cell types. That is, pre-meiotic and meiotic female germ cells showed reduced sequence conservation and expression of younger genes on par with their male counterparts. We also found evidence of a highly permissive chromatin landscape and promiscuous transcription in female meiotic germ cells, as previously described for male meiotic germ cells^10–12^. These observations suggest that the “out of the testis” hypothesis^4–9^, which proposes new genes often evolve in the testis because of a permissive chromatin environment, can be extended to the ovaries. During spermatogenesis, the promiscuous transcription is accompanied by the expression of unique splicing variants^11,88^ and translational buffering^9,11^. Future studies using long-read sequencing and ribosome profiling could determine whether these phenomena also occur in developing female germ cells.

## Methods

### Biological sample collection and ethics statement

**Human Samples:** Human prenatal gonad samples were provided by the MRC-Wellcome Trust Human Developmental Biology Resource (HDBR) and obtained from elective pregnancy terminations. Donors made the decision to contribute tissue voluntarily, following counselling regarding pregnancy termination, and provided written informed consent for the collection of fetal material. All prenatal samples, except one identified as XXY (Klinefelter syndrome), had normal karyotypes and were classified according to a particular Carnegie stage or week post conception (WPC) based on their external physical appearance and measurements. Samples were collected at weeks 5-7, 8, 10-12, 17, and 19-21 (Supplementary Table 1), covering the processes of sex determination and sex-specific differentiation of major cell types. The use of human samples was approved by an ERC Ethics Screening panel (associated with H.K.’s ERC Consolidator Grant 615253, OntoTransEvol) and ethics committees in Heidelberg (authorization S-220/2017), North East-Newcastle & North Tyneside (REC reference 95 18/NE/0290), and London-Fulham (REC reference 18/LO/0822).

**Marmoset Samples:** Animal experiments for obtaining marmoset embryos were conducted in accordance with German legislation and approved by the Lower Saxony State Office for Consumer Protection and Food Safety (Niedersächsisches Landesamt für Verbraucherschutz und Lebensmittelsicherheit, LAVES) ethics committee under license number 425020412/0708. Embryos at specific developmental stages, approximately corresponding to human Carnegie stages 16 to 23 (gestational day (GD) 74 to 95)^89,90^ were collected via hysterotomy as previously described^91^. A specialized veterinarian performed all surgical procedures, including anesthesia and analgesia. Additionally, postnatal samples from newborn and 3weeks-old marmoset were collected following euthanasia for reasons unrelated to participation in this study. Gonadal samples were dissected, snap-frozen in liquid nitrogen and stored at −70°C.

### Nuclei Preparation

Nuclei were extracted as previously described^92^. Briefly, frozen tissue was homogenized on ice in a buffer containing 250 mM sucrose, 25 mM KCl, 5 mM MgCl2, 10 mM Tris-HCl (pH 8), 0.1% IGEPAL, 1 µM DTT, 0.4 U/µl Murine RNase Inhibitor (New England BioLabs), 0.2 U/µl SUPERase-In (Thermo Fisher Scientific), cOmplete Protease Inhibitor Cocktail (Roche). The tissue was disrupted by trituration and/or using a micropestle. After a brief incubation, unlysed tissue debris was removed by low-speed centrifugation (100g for 1 minute at 4°C). The supernatant was then centrifuged at 400g for 4 minutes to separate the nuclei (pellet) from the cytoplasmic fraction (supernatant). Nuclei were washed once or twice in the homogenization buffer and resuspended in a storage buffer containing 430 mM sucrose, 70 mM KCl, 2 mM MgCl2, 10 mM Tris-HCl (pH 8), 0.4 134 U/µl Murine RNase Inhibitor, 0.2 U/µl SUPERase-In, cOmplete Protease Inhibitor Cocktail. If needed, the nuclei were filtered using 40 µm Flowmi strainers. Nuclei concentration was determined by staining with Hoechst DNA dye or propidium iodide and counting on a Countess II FL Automated Cell Counter (Thermo Fisher Scientific). Each library, snRNA-seq or snATAC-seq, was generated using approximately 15,000 nuclei. For early marmoset gonad samples (GD74 and GD80), around 5000 nuclei were used for library generation due to the small size of the gonads at these stages.

### RNA extraction and sample quality control

RNA was extracted from the cytoplasm extracts or nuclei suspensions by mixing them with RLT buffer (supplemented with 40 mM DTT) and 100% ethanol in a 2:7:5 ratio. The RNA was then purified using the RNeasy Micro Kit from Qiagen. The quality of the extracted RNA was assessed using the Fragment Analyzer (Agilent; RRID:SCR_019417), and all samples had RNA quality numbers (RQN) above 8.

### Library preparation and sequencing

**snRNA-seq:** Single-cell barcoding and library preparation were performed using Chromium Single Cell 3’ Reagent Kits (v3 chemistry) and the Chromium Controller instrument (10X Genomics; RRID:SCR_019326), following the manufacturer’s protocols. The cDNA was amplified using 12 PCR cycles. The Qubit Fluorometer (Thermo Fisher Scientific; RRID:SCR_018095) was used to quantify the libraries, and the average fragment size was assessed using the Fragment Analyzer. Sequencing of the libraries was carried out on Illumina NextSeq 550 (RRID:SCR_016381; 28 cycles for Read 1, 8 cycles for i7 index, 56 cycles for Read 2).

**snATAC-seq:** Nuclei were processed using Chromium Single Cell ATAC Reagent kits (v1) and the Chromium Controller instrument (10x Genomics; RRID:SCR_019326), following the manufacturer’s guidelines. The process included tagmentation, single cell barcoding, and library preparation. The libraries were then amplified through 10 PCR cycles and quantified using a Qubit Fluorometer (Thermo Fisher Scientific; RRID:SCR_018095). The average fragment size of the libraries was determined using a Fragment Analyzer (Agilent; RRID:SCR_019417). Sequencing was performed on Illumina NextSeq 550 (RRID:SCR_016381) with 34 cycles for both Read 1 and Read 2, 8 cycles for the i7 index, and 16 cycles for the i5 index.

### snRNA-seq data processing and quality control

For each sample, we demultiplexed the raw sequencing data using bcl2fastq. We aligned the reads to the human genome (GRCh38) or the marmoset genome (CalJac4) and counted the unique molecular identifiers (UMIs) with the 10X Genomics Cell Ranger (version 4.0.0) count pipeline, retaining intronic reads. To identify valid barcodes stemming from droplets containing a nucleus (as opposed to empty droplets containing only ambient RNA), we clustered the barcodes by their number of reads and the fraction of intronic UMIs with a Bayesian Gaussian mixture model with a Dirichlet process prior (using the function *BayesianGaussianMixture* from scikit-learn 0.20.1, python 3.6.6). We choose the cluster with its centroid having the highest fraction of intronic UMIs and the largest number of reads, as droplets containing a nucleus have a higher fraction of intronic UMIs and a larger number of reads than empty droplets with ambient RNA (stemming mostly from the cytoplasm of broken cells). We then filtered out barcodes containing more than one nucleus using Scrublet^93^ (version 0.2, python 3.6.6) by keeping only those with a doublet score less than 0.5. Furthermore, we removed nuclei with less than 300 unique genes expressed and less than 20,000 UMIs. After these filtering steps, we retained 69,597 human testis, 57,774 human ovary, 9,612 marmoset testis and 35,491 marmoset ovary nuclei, as well as 8,936 XXY human testis nuclei of high quality.

### snRNA-seq dataset integration and normalization

We removed batch effects and integrated across stages within each organ and species using the rliger package (version 1.0.1, R 4.1.3)^19^. We used default parameters for normalization, scaling, and selection of highly variable genes. We removed cell cycle, ribosomal and mitochondrial genes from the variable features to remove integration bias and performed integrative non-negative matrix factorization using the *optimizeALS* function with parameters k = 200, lambda 30 and 3 repetitions, followed by *quantile_norm* with knn_k = 100, min_cells = 200 and quantiles = 100. We generated integrations for each sex and species separately, and complete integrations for each species, including both, XX and XY cells.

### snRNA-seq dimensionality reduction, clustering and annotation

We performed dimensionality reduction using Uniform Manifold Approximation and Projection^94^ (UMAP) with n_neighbors = 100 and clustered the datasets using the *louvainCluster* function with resolution = 12 of rliger, which internally uses the Seurat Louvain clustering implementation. We converted the rliger object to a Seurat object for further analysis using the *ligerToSeurat* function.

We identified the main cell types based on the expression of known human marker genes and their 1:1 orthologs in marmoset. See Extended Data Figs. 1-4 and Supplementary Table 2 for the marker genes used to annotate the cells.

### Calculation of X:A ratio

We estimated the fraction of biallelic expression in our data using a modified version of the *scanForHeterozygotes* function of the AllelicImbalance^95^ R package, which we adapted to work with the large high dimensional matrices of single cell data. In short, the function takes reads aligned to the genome from all cells of a sample and measures the frequency of alternative bases at each position, requiring a minimum of 20 reads, a maximum major allele frequency of 0.9, a minimum minor allele frequency of 0.1 and a minimum biallelic frequency of 0.9 to identify major and minor alleles for each position. We then count for each cell the number of reads supporting either the minor or major allele and recorded the fraction of positions with reads supporting both alleles. We ran this analysis on all our snRNA-seq and snATAC-seq datasets which included at least 500 germ cells.

We calculated the X:A expression ratio using pseudobulk replicates. We first selected broadly expressed genes to reduce bias introduced by cell type specific gene expression, an approach similar to the one used previously^35^. For this, we created pseudobulks for each broad cell type lineage using the *AggregateExpression* function in Seurat on the human ovary and testis snRNA-seq dataset, down sampled to the number of cells in the smallest group (n = 754) and selected genes with a minimum count of greater than 1, giving us a set of 572 X-linked and 14964 autosomal genes. We then created pseudobulks for each cell type, sex, and replicate, grouping all somatic cell types together and using *AggregateExpression* on the dataset, downsampled to the number of cells in the smallest group (n = 806). Finally, we calculated the X:A ratio by dividing the mean expression (CPM) of x-linked genes by the mean expression of autosomal genes.

### Reanalysis of published XCI data

To reanalyse the data published by Chitiashvili et al.^15^, we downloaded the cell by gene matrices for the female samples and performed data integration following the Seurat workflow (*SCTransform*, *FindIntegrationAnchors*, *IntegrateData*, *RunPCA*, *RunUMAP*) with default parameters. We annotated the germ cell states based on markers and annotated the remaining cells as “somatic”. We then calculated the X:A ratio by following the same protocol as for our data (Extended Data Fig. 6a).

The correct estimation of the X:A ratio depends on the selection of genes, specifically ubiquitously expressed genes^35^. Additionally, calculating the X:A ratio per cell can be skewed by random technical dropouts and comparing all cells of multiple clusters with each other loses the information of biological variability captured by multiple replicates, which is why in our analysis, we chose to estimate the X:A ratio based on pseudobulks from biological replicates. Including all (protein-coding and non-coding) genes, including those with restricted or cell type-specific expression, and calculating the X:A ratio based on individual cells (using the per cell sums of SCTransformed^96^ or log-normalized and scaled expression values of X-linked and autosomal genes), recapitulates the results of Chitiashvili et al., showing a relatively low X:A ratio for PGCs and a higher X:A ratio in meiotic oogonia (Extended Data Fig. 6b). Because the authors of that study did not disclose details on how they calculated the X:A ratio, we can only assume that the conflicting findings can be traced back to this calculation.

### snATAC-seq data processing and quality control

For each sample, we demultiplexed the raw sequencing data using the CellRanger-ATAC (version 1.2.0) mkfastq pipeline, which internally uses bcl2fastq. We aligned the reads to the human genome (GRCh38) or the marmoset genome (CalJac4) with the Cell Ranger-ATAC count pipeline. We imported the aligned reads into R (version 4.1.3) using the ArchR package (version 1.0.2) and retained cells with a minimum transcription start site enrichment score of 3 and with at least 5,000 and at most 10^5^ mapped ATAC-seq fragments. We excluded fragments of less than 10bp and more than 2,000 bp in size. We then removed potential doublets by filtering out cells with more than 45,000 fragments or with a doublet score of higher than 4, as calculated by the *addDoubletScores* function of ArchR. After filtering, we retained 51,966 human testis, 58,386 human ovary, 12,185 marmoset testis and 12,294 marmoset ovary nuclei, as well as 11,022 human XXY testis nuclei of high quality.

### snATAC-seq dimensionality reduction and clustering

We first used iterative latent semantic indexing (LSI) implemented in the ArchR function *addIterativeLSI* with 5 iterations with increasing clustering resolution (0.1, 0.2, 0.4, and 0.8), sampling 20,000 cells each time and using the 100,000 most variable features to reduce the dimensionality. Based on the resulting components, we then performed Louvain clustering and generated a UMAP embedding using the *addClusters* (resolution = 1.5) and *addUMAP* (minDist = 0.25, metric = “correlation”, nNeighbors = 100) functions of ArchR, respectively.

### snATAC-seq label transfer and annotation

We integrated our snATAC-seq cells with our previously annotated snRNA-seq cells using the *addGeneIntegrationMatrix* function in ArchR, which internally uses the canonical correlation analysis (CCA) implementation of the Seurat package^97^ to find the most similar snRNA-seq cell for each snATAC-seq cell based on the gene expression and gene score (calculated from accessibility within the gene body and putative regulatory elements), respectively. We constrained the integration to matching developmental stages, or the closest match when no exact match was available. We then used the detailed snRNA-seq annotation to transfer the labels to the corresponding snATAC-seq cells and confirmed this annotation by plotting the gene score of cell type marker genes in a UMAP embedding.

### Annotation of snATAC-seq peaks

We identified reproducible and robust regions of accessible chromatin following a protocol previously established in our lab^64^ using the *addGroupCoverages* and *addReproduciblePeakSet* functions of the ArchR package, which internally called peaks using MACS2^98^ for each cell state and assessed reproducibility based on our biological (or technical) replicates, requiring a peak to be called in at least two replicates to be considered reproducible. We further filtered our peaks by requiring them to be present in at least 5% of at least one cell state to be considered robust. We added motif annotations using the ArchR function *addMotifAnnotations* based on the cisBP motif set^99^. Using the ArchR implementation of chromVAR^100^ (*addBgdPeaks* and *addDeviationsMatrix*), we inferred per-cell TF activity. The peaks were annotated as either promoters, exonic, intronic or intergenic by ArchR based on a −2000/+100 bp region around the peaks. We supplemented this annotation using a previously published data set on human lncRNAs^8^.

We assigned peaks to putative target genes by first integrating our snRNA-seq data with our snATAC-seq data using the *addGeneIntegrationMatrix* function of ArchR which is based on the *FindTransferAnchors* function of Seurat^97^. We chose an integration approach constrained by developmental stage and sex, choosing the closest available stage in cases where the snATAC-seq and snRNA-seq data were not exactly matched (integrating human CS20 snATAC-seq data with CS22 snRNA-seq data, and marmoset GD92 snATAC-seq data with GD95 snRNA-seq data). We then used the ArchR function *addPeak2GeneLinks* to correlate peak accessibility with gene expression, employing at first no correlation cutoff. We filtered the peak to gene linkages using an empirically established cutoff using correlations within a 250kb window and across different chromosomes, giving us a predicted FDR of 0.01 (r = 0.37 in human, r = 0.44 in marmoset). We then annotated the peaks based on their associated gene into four biotypes – protein-coding, lncRNA, small RNA and other – using Ensembl version 102.

### Global changes in chromatin landscape

To identify regions of nucleosome-depleted chromatin that are shared between the sexes or sex-specific in our human snATAC-seq data, we first matched corresponding male and female cell types. For the supporting lineage, we included coelomic epithelial cells and common somatic progenitor cells, which are present in our testis and ovary samples, as well as developing Sertoli cells, pregranulosa cells and developing granulosa cells. We split the cells into four developmental phases – CS17-22, 10-12WPC, 17-19WPC and 20-21WPC. For the germ cell lineage, we split the cells by their progression through germ cell development (PGCs, FGCs, and pre-spermatogonia/pre-meiotic oogonia), rather than the sampled time points, as germ cells develop asynchronously. We then down sampled the cells from the corresponding male and female groups to the same number and assessed whether a peak was sex-specific or shared with 1000 bootstrap replicates. We defined a peak as male-specific if it was present in at least 4% of male cells and where the percentage of male cells with this peak is at least twice as high as that of female cells – and vice versa for female-specific peaks. We defined a peak as shared, when it was present in at least 4% of male and female cells, and the absolute log2 fold-change between the percentage of male cells with the peak and female cells with the peak was below 1.

In supporting cells, we observed similar ratios of sex-specific and shared CREs between the autosomes and the X chromosome (Extended Data Fig. 7b). However, in germ cells, there are considerably more female-specific CREs on the X chromosome (Extended Data Fig. 7c), likely due to the increased peak detection power from the two active X chromosomes during XCR in female germ cells. Therefore, we excluded the X chromosome from further analyses in this section if not specified otherwise.

### Dating of cis-regulatory regions

We assigned a minimum evolutionary age to each peak in our human dataset by aligning their sequences to the genomes of 18 vertebrate species with increasing phylogenetic distances (Supplementary Table 8) downloaded from UCSC. We used liftover with the parameters: minMatch = 0.1, minSizeQ = 50 and minSizeT = 50 to assess the presence of the sequences in other species. Using estimated divergence times between human and the respective species from TimeTree (http://www.timetree.org/)^101^, we assigned minimum ages to all peaks depending on the species with the largest divergence time in which the sequence was still found.

### Gene regulatory networks

We inferred gene regulatory networks using the CellOracle python package (version 0.10.1, python version 3.8). For this, we built a base GRN based on our human germ cell and somatic snATAC-seq data separately using the default motif set for vertebrates (gimme.vertebrate.v5.0) included with CellOracle. We then used our snRNA-seq data (converted from Seurat to AnnData^102^ version 0.8.0) to build sex and cell state specific GRNs following the CellOracle documentation. We removed weak network edges using a *P* value filter of 0.001 and by choosing the top 5000 edges ranked by edge strength.

### Pseudotime and trajectory changes

To create pseudotime trajectories recapitulating cell type differentiation, we first separated germ cells and somatic cells. For the somatic trajectories, we included all cell types of the major four lineages, (Sertoli, Leydig, granulosa, and ovarian stroma), coelomic epithelial cells, and common somatic progenitor cells in a merged Seurat^97^ object. We then used Monocle 3^70–72^ to predict a branching pseudotime trajectory starting at the coelomic epithelial cells of the earliest sample. We smoothed the gene expression along the four resulting trajectories using a negative binomial generalized additive model using the tradeSeq package^103^. We used the associationTest function of the tradeSeq package to identify genes that show a dynamic expression pattern along one of the four pseudotime trajectories. For the female and male germ cells, we included all germ cell states and smoothed the gene expression along the two trajectories using a negative binomial generalized additive model and performed association tests using tradeSeq. To cluster genes, we used hierarchical clustering on Spearman distance with the Ward d2 method.

To identify genes with conserved or changed trajectories between human, marmoset, and mouse, we first integrated our human and marmoset datasets with published mouse scRNA-seq datasets for germ cells from embryonic day (E) 12.5 through E16.5^76^ and somatic cells from E10.5 through E16.5^77^ separately for each lineage (germ cells, supporting cells, steroidogenic cells). For this, we limited our genes to 1:1 orthologs between all three species and integrated the datasets using the *IntegrateData* function of Seurat^97^ (version 4.3.0.1), followed by dimensionality reduction using PCA and UMAP. Using this joint embedding, we ordered the cells along a pseudotime trajectory following the development of the cell types using Monocle 3^70–72^ (version 1.3.1). We created 10 equal bins along the pseudotime trajectory and created pseudobulks for each replicate using the *AggregateExpression* function in Seurat^97^. We selected dynamically expressed genes by measuring the distance correlation of the gene expression with the pseudotime bins and setting a threshold of 0.3. We then calculated the distance correlation, Spearman correlation, and root mean square error (RMSE) of the gene expression along the bins pairwise between the three species for each gene. Genes with a distance correlation of less than 0.3 between the species were classified as “undefined” to filter out noisy gene expression trajectories. We set the following thresholds for genes with conserved trajectories: distance correlation > 0.5, Spearman correlation > 0, and RMSE > 0.4. We classified genes where one of the species had values below these thresholds as different in that species, while classifying genes where all species showed values below these thresholds as “undefined”.

### Estimation of the evolutionary rate of change

To estimate the mode and strength of evolution of genes in the human gonadal cell types, we downloaded human-to-mouse homolog dN and dS data from Ensembl (version 99) for all one-to-one orthologs. We then calculated the mean dN/dS and the 95 percent confidence interval for the marker genes of the cell states. For our marmoset data, we calculated the dN/dS values in the same way using marmoset-to-mouse homolog dN and dS data from Ensembl (version 99). We also calculated the dN/dS for published mouse data (germ cell data from Mayère et al, 2021^76^ and somatic cell data from Mayère et al, 2022^77^) using mouse-to-human homolog dN and dS data from Ensembl (version 99). To estimate the mean phylogenetic age of genes expressed in each cell state of our human data, we used the GenTree^81,82^ resource (http://gentree.ioz.ac.cn/) which assigns a higher score to newer genes based on the branch leading to humans on which the gene could first be detected. We calculated the average and 95 percent confidence interval of the GenTree scores of the marker genes for each cell state.

For our human snATAC-seq datasets, we used phastCons^80^ and phyloP^104^ scores based on multiple alignments of 100 vertebrate species (downloaded from UCSC) and 241 mammalian species (from the Zoonomia project^105^), respectively, to assess sequence constraint of the identified peaks. For this, we used bigWigAverageOverBed^106^ tool on sliding windows of 100bp width and chose the mean phastCons and phyloP scores of the most conserved window within each peak. To assess the conservation of CRE activity, we used liftover^107^ to identify homologous sequences of accessible (CPM ≥ 5) human CREs in marmoset. We split the peaks into peaks accessible in germ cells or somatic cells, and in early accessible (10-12WPC in human, GD92-95 in marmoset) and late accessible (19-21WPC in human, NB in marmoset). We then measured the fraction of homologous sequences that were accessible in both species.

### Tissue specificity

We used time and tissue specificity scores (tau)^108^, previously calculated for the ovary and testis^5^. The time specificity scores were calculated across developmental time points in the ovary and testis, while the tissue specificity scores were determined across tissues (including brain, heart, kidney, liver, and gonads). These scores range from 0 to 1, where 0 indicates low and 1 indicates high specificity.

### XCI escapees

We used a list of reported human XCI escaping genes published by Tukiainen et al.^109^.

### DSD and abnormal fertility gene lists

We retrieved lists of genes that are reportedly associated with human DSDs from previously published work^65–67^ and downloaded a list of genes that are associated with abnormal fertility/fecundity in mice using the identifier MP:0002161 from the IMPC database (www.mousephenotype.org)^110^.

### Human sex-related pathogenic variants

We retrieved the ClinVar (https://www.ncbi.nlm.nih.gov/clinvar/)^111^ variant file (GRCh38, version 20240716) and filtered it for pathogenic or likely pathogenic variants and for conditions including any of the terms ovary/ovarian, testis/testicular, sperm, oogonia/oocytes, fertility, fecundity.

### Phylogenetic trees

Divergence times and phylogenetic trees in this study are based on TimeTree (http://www.timetree.org/)^101^.

### GO term enrichment analyses

We performed enrichment analyses of Gene Ontology^61,62^ terms using the g:Profiler^112^ R package.

### General statistics and visualizations

Unless otherwise specified, we used R version 4.2.2 installed in a custom conda environment on an Ubuntu 22.04.4 LTS system to analyse and create plots.

## Data and code availability

The datasets created for this study are available at ArrayExpress under the following accessions: E-MTAB-15021 (human snRNA-seq data, reviewer access: https://www.ebi.ac.uk/biostudies/ArrayExpress/studies/E-MTAB-15021?key=d7eccd44-4493-4875-b9c9-b0fe882c64c5), E-MTAB-15028 (human snATAC-seq data, reviewer access: https://www.ebi.ac.uk/biostudies/ArrayExpress/studies/E-MTAB-15028?key=90bc4afb-d8dc-4a7f-ae60-0765fb542bcc), E-MTAB-15025 (marmoset snRNA-seq data, reviewer access: https://www.ebi.ac.uk/biostudies/ArrayExpress/studies/E-MTAB-15025?key=7519c8b7-2ad6-4234-b97f-b2019a58378a) and E-MTAB-15024 (marmoset snATAC-seq data, reviewer access: https://www.ebi.ac.uk/biostudies/ArrayExpress/studies/E-MTAB-15024?key=e11e18cd-0000-4f72-b5e9-68d241a52a85).

We provide an interactive web application to explore the processed data, hosted at https://apps.kaessmannlab.org/primate_gonadogenesis/. Custom code used for the analyses in this study is available at https://gitlab.com/kaessmannlab/primate_gonadogenesis.

## Acknowledgements

We acknowledge all members of the Kaessmann lab for discussions, K. Hall for assistance, Dr. Sophie Mißbach, Dr. Michael Heistermann (DPZ Laboratory for Endocrinology), Angelina Berenson, Ulrike Goedecke, Nicole Umland and the animal caretakers of the DPZ animal facility. We acknowledge the Joint MRC/Wellcome (MR/R006237/1) Human Developmental Biology Resource for providing human samples. We acknowledge support by the state of Baden-Württemberg through bwHPC and the German Research Foundation (DFG) through grant INST 35/1597-1 FUGG. We acknowledge the data storage service SDS@hd supported by the Ministry of Science, Research and the Arts Baden-Württemberg (MWK) and the German Research Foundation (DFG) through grant INST 35/1503-1 FUGG. The purchase of the NextSeq 550 instrument was supported by the Klaus Tschira Foundation. This research was supported by grants from the ERC (grant no. 615253, OntoTransEvol) and the German Research Council (DFG, grant no. SFB 873) to H.K.

## Author contributions

N.T., A.F. and H.K. conceived and organised the study. A.F. performed experiments with support from C.S., J.S., N.M. and R.F and guidance from M.S. N.T. and A.F. analysed data with support from I.S. and F.M. E.W. collected marmoset samples, C.D. performed marmoset experimentation and R.B. provided marmoset samples. S.L. provided human samples. I.S., N.M., M.S., F.M. and M.C.-M. provided critical discussions. N.T. developed the web-application. H.K. provided funding. H.K. and R.B. supervised the study. N.T. and A.F. drafted the manuscript with critical review by H.K. All authors provided feedback on drafts and approved its final version.

## Extended Data Figures

**Extended Data Fig. 1:**
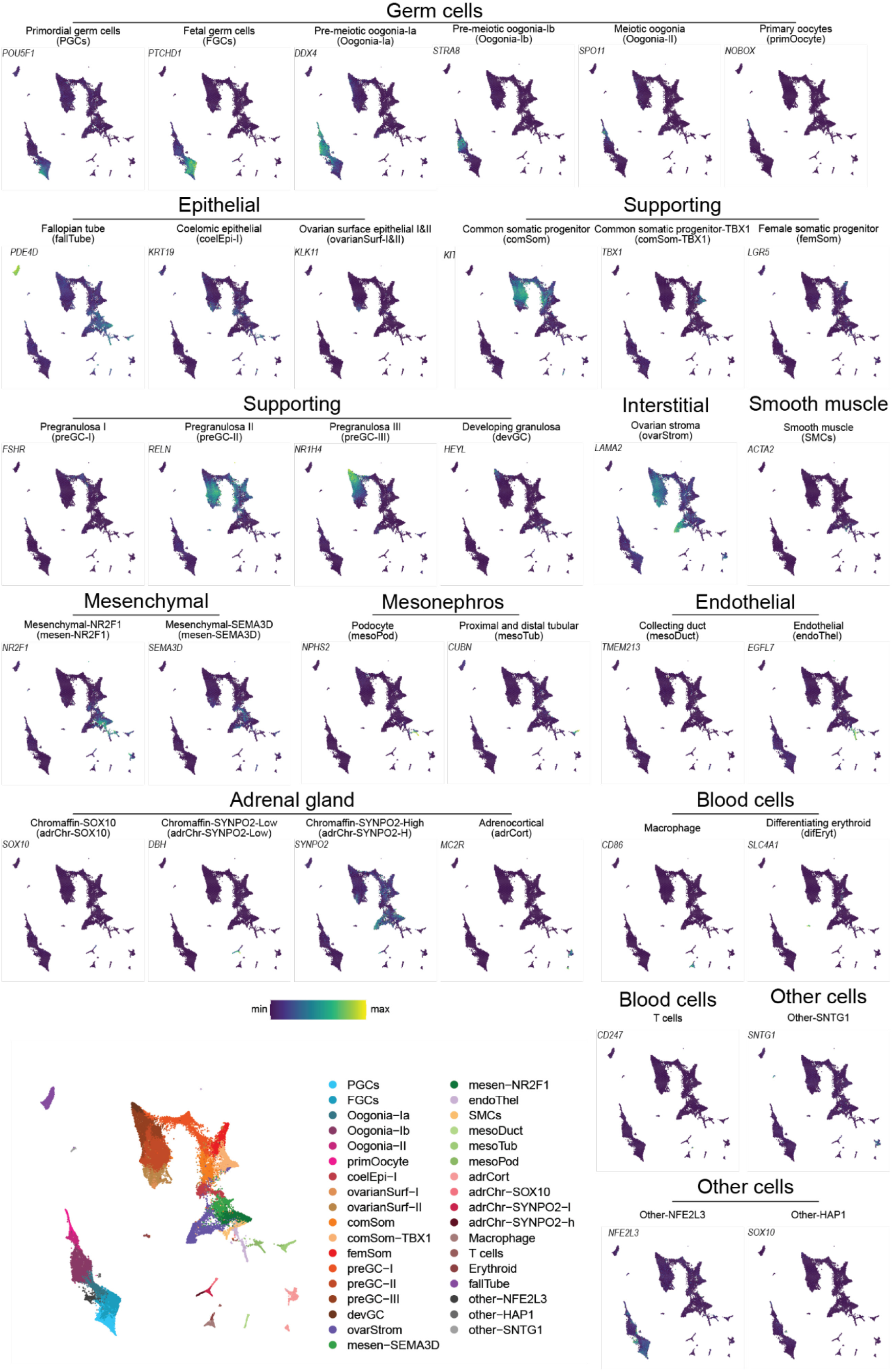
Human female gonad marker genes. UMAP embeddings of the human ovary

**Extended Data Fig. 2:**
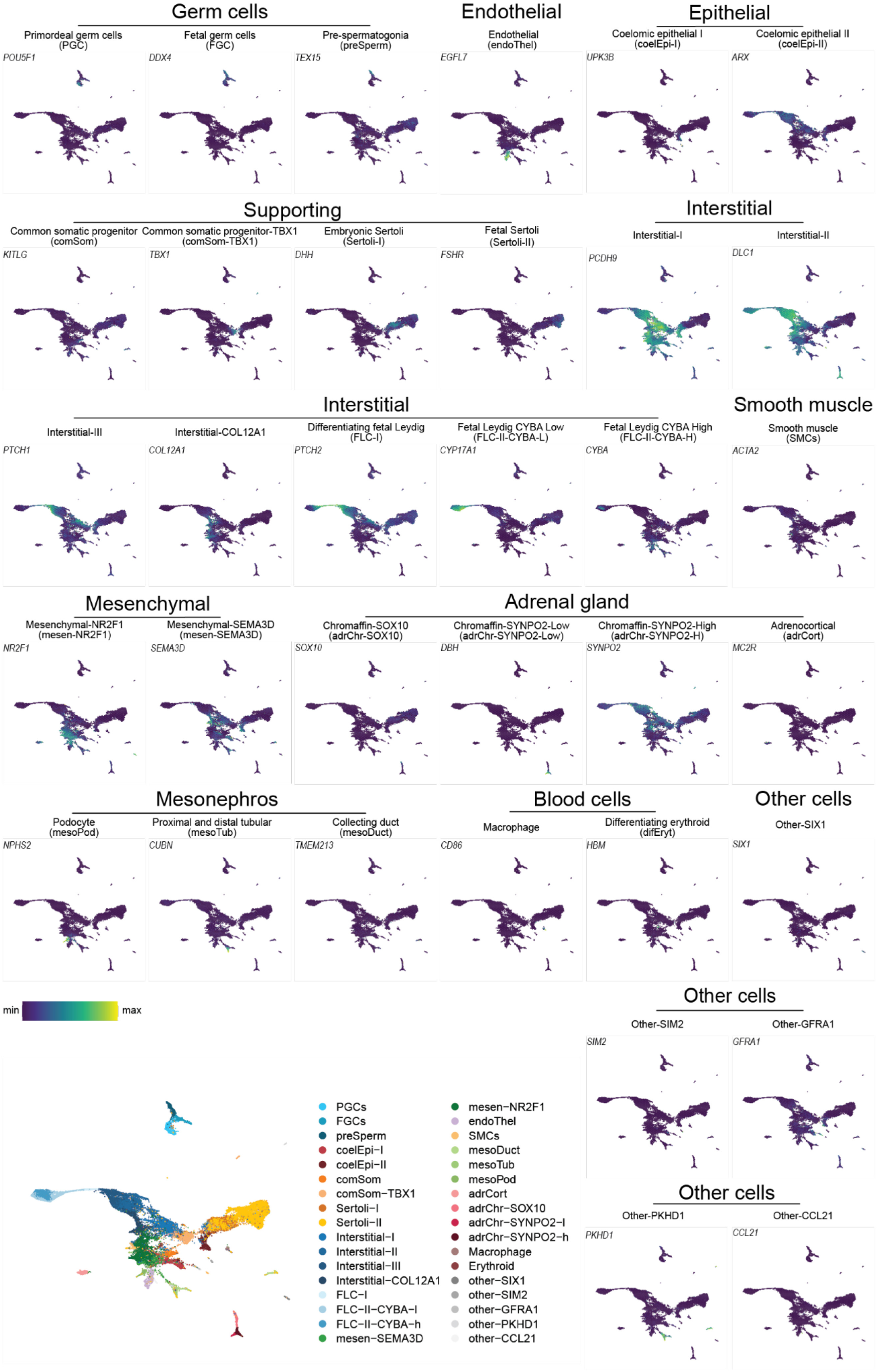
Human male gonad marker genes. UMAP embeddings of the human testis

**Extended Data Fig. 3:**
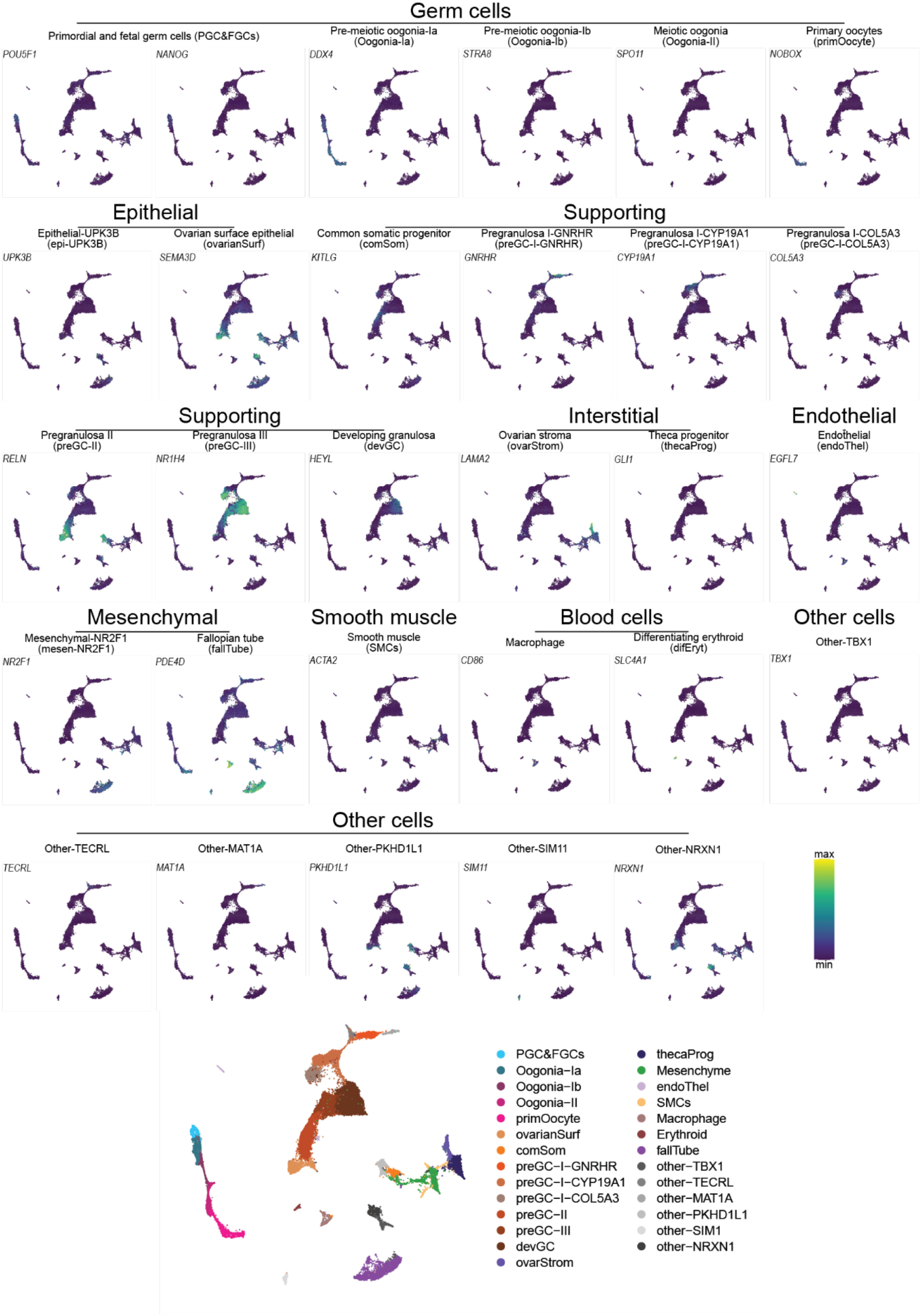
Marmoset female gonad marker genes. UMAP embeddings of the marmoset ovary snRNA-seq dataset coloured by the expression of marker genes for each cell type.

**Extended Data Fig. 4:**
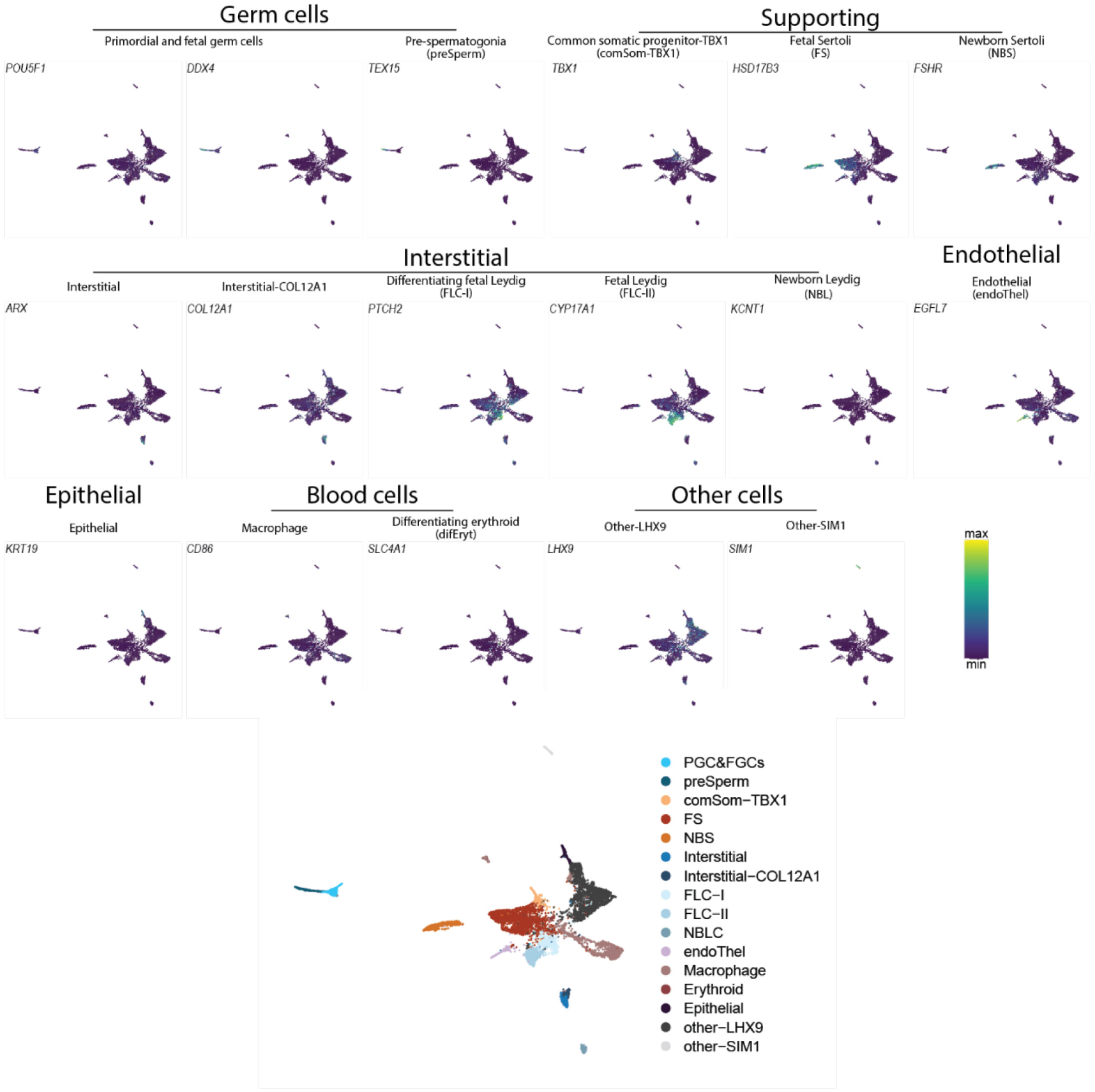
Marmoset male gonad marker genes. UMAP embeddings of the marmoset testis dataset snRNA-seq coloured by the expression of marker genes for each cell type.

**Extended Data Fig. 5:**
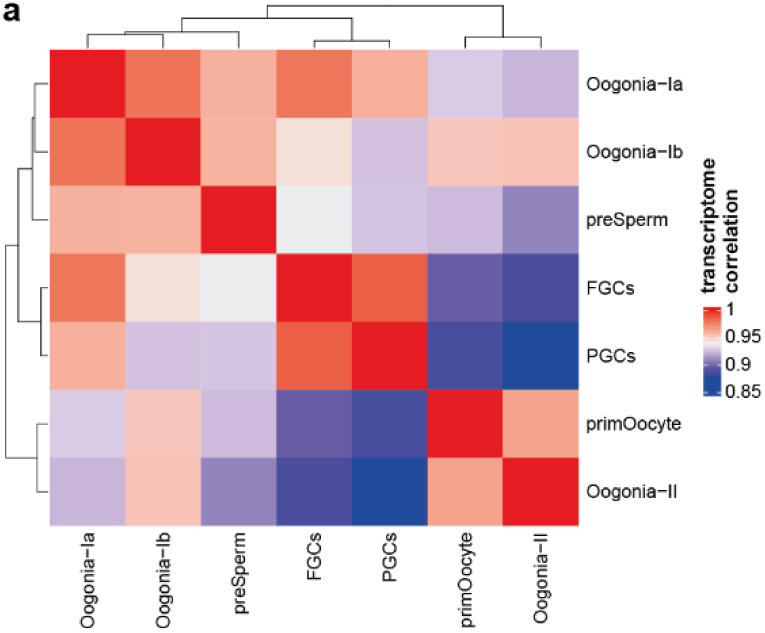
Transcriptome similarities. Heatmap of Spearman’s correlation coefficients between snRNA-seq pseudobulks of germ cell states.

**Extended Data Fig. 6:**
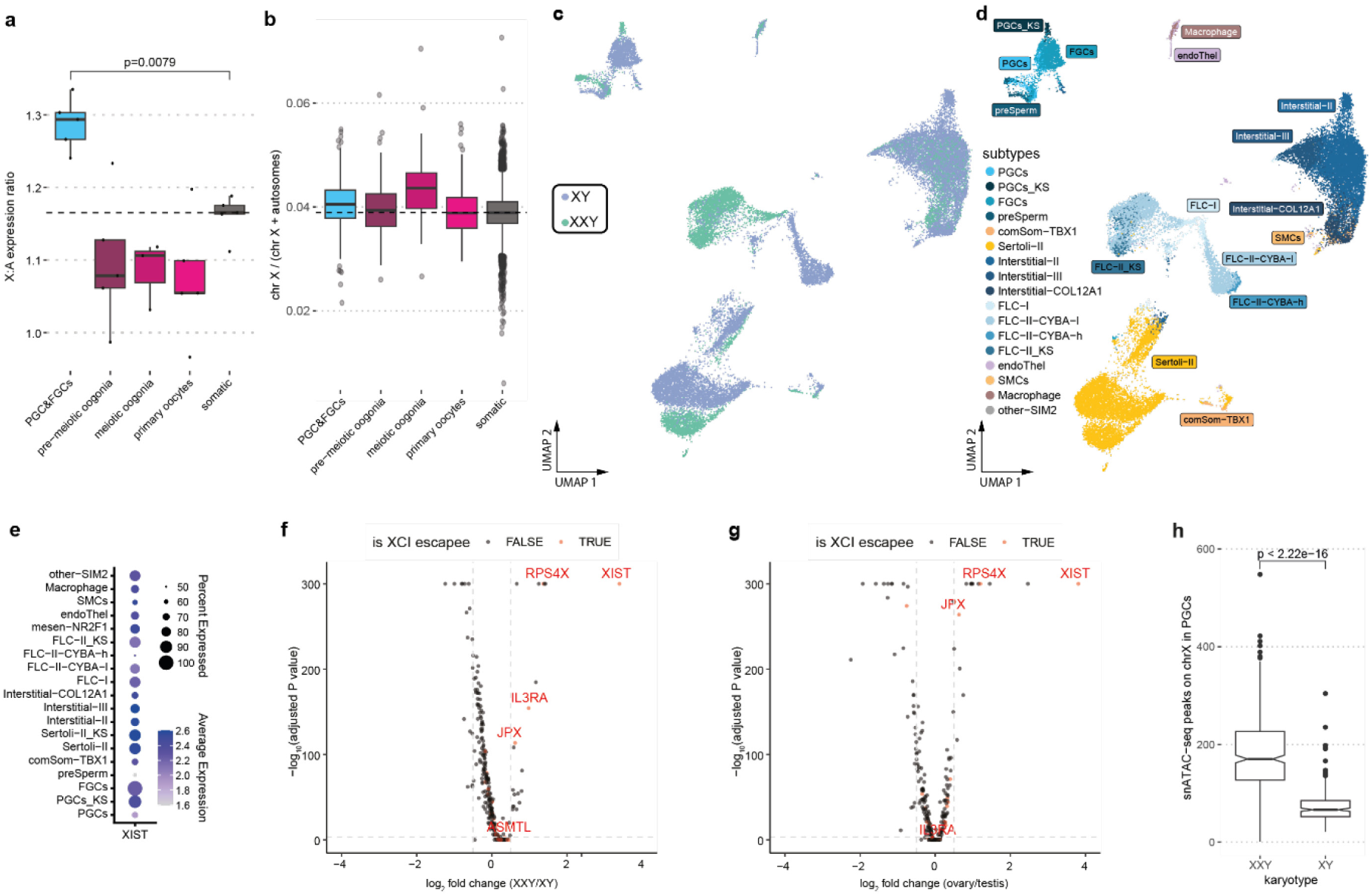
XCI and Klinefelter syndrome. **a,b** Box plots of X to autosome expression ratios in human ovary data from Chitiashvili et al.^15^ calculated using our method (**a**; dots indicate pseudobulk replicates; see Methods) and as described in their methods (**b**; dots represent individual cells). **c**,**d**, UMAP embeddings of snATAC-seq datasets of 12WPC XY and 13WPC XXY testis coloured by karyotype (**c**) and (**d**) cell type. **e**, XIST expression in XXY testis cell types. **f**, Volcano plot of differentially expressed X-linked genes between somatic cells of 13WPC XXY and 12WPC XY testes. A larger than 0 log_2_ fold change indicates higher expression in XXY than in XY. Red points denote known XCI escaping genes. **g**, Volcano plot of differentially expressed X-linked genes between somatic cells of 12WPC ovaries and testes. A larger than 0 log_2_ fold change indicates higher expression in ovaries than in testes. **h**, snATAC-seq peaks on the X chromosome in XXY and XY PGCs. The *P* value was calculated using the Wilcoxon rank sum test. In **a**, **b** and **h**, boxes represent the first to third quartiles and whiskers extend to 1.5 times the inter-quartile range from the box hinges. The horizontal lines mark the median.

**Extended Data Fig. 7:**
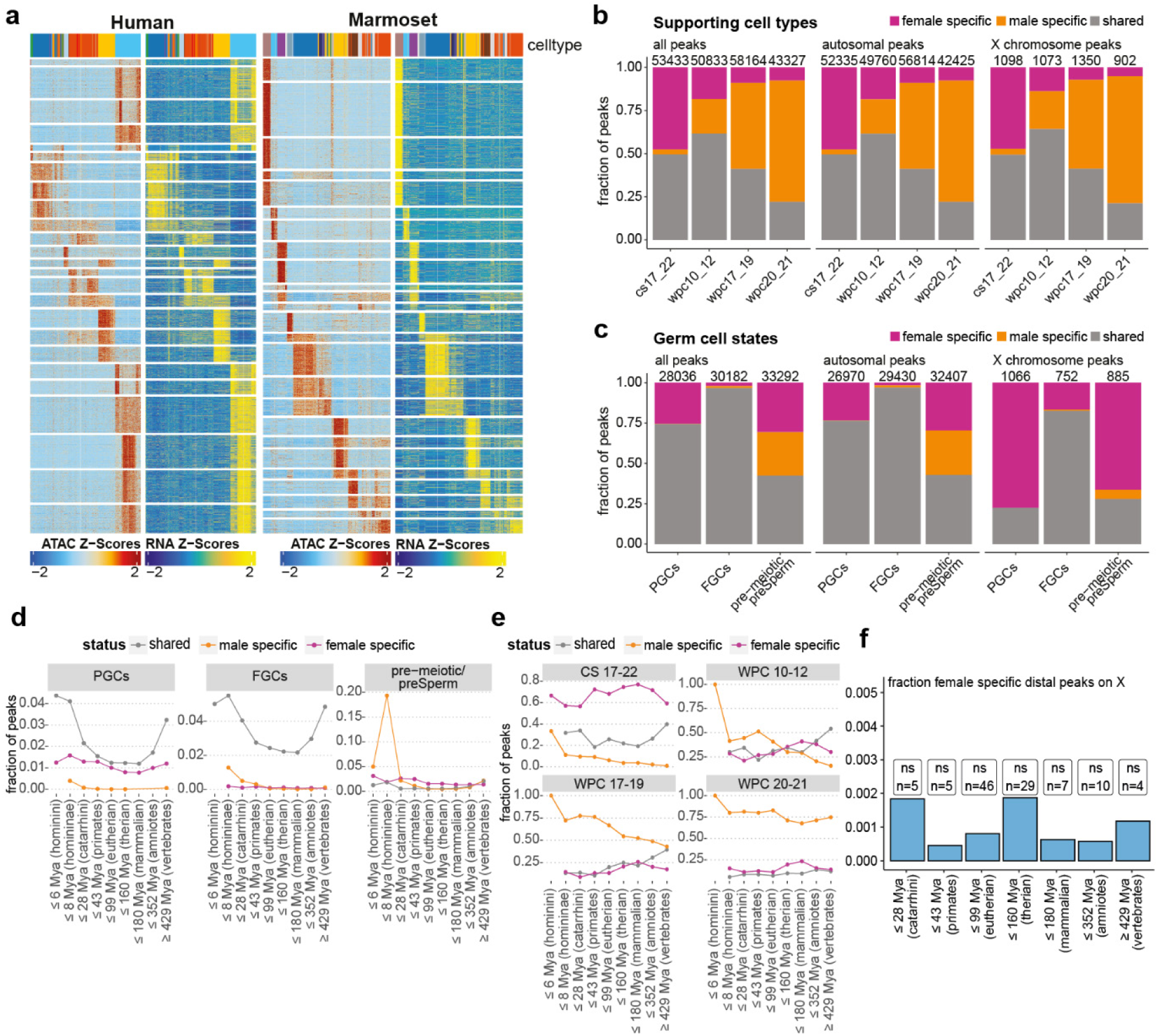
Chromatin dynamics. **a,** Heatmap showing snATAC-seq gene scores and snRNA-seq gene expression z-scores across human and marmoset gonadal cell types (see Extended Data Fig.1,2,3 and 4 for colour legends of cell types). **b**,**c**, Fractions of shared and sex-specific peaks per stage in human supporting cells (**b**) and per cell state in human germ cells (**c**). Sub-panels show the fractions for peaks on all chromosomes, only autosomes, and only the X chromosome. Numbers above the bars show the total number of shared and sex-specific peaks per stage or cell state. **d**,**e**, Fractions of human shared and sex-specific autosomal CREs in each age group in germ cells (**d**) and supporting cells (**e**). **f**, Fraction of human female-specific intergenic peaks in supporting cells on the X chromosome in each age group. Labels show the significance (Fisher’s exact test) of the overrepresentation of female-specific peaks on the X chromosome (ns = p>0.05, Benjamini-Hochberg adjusted). Mya, million years ago.

**Extended Data Fig. 8:**
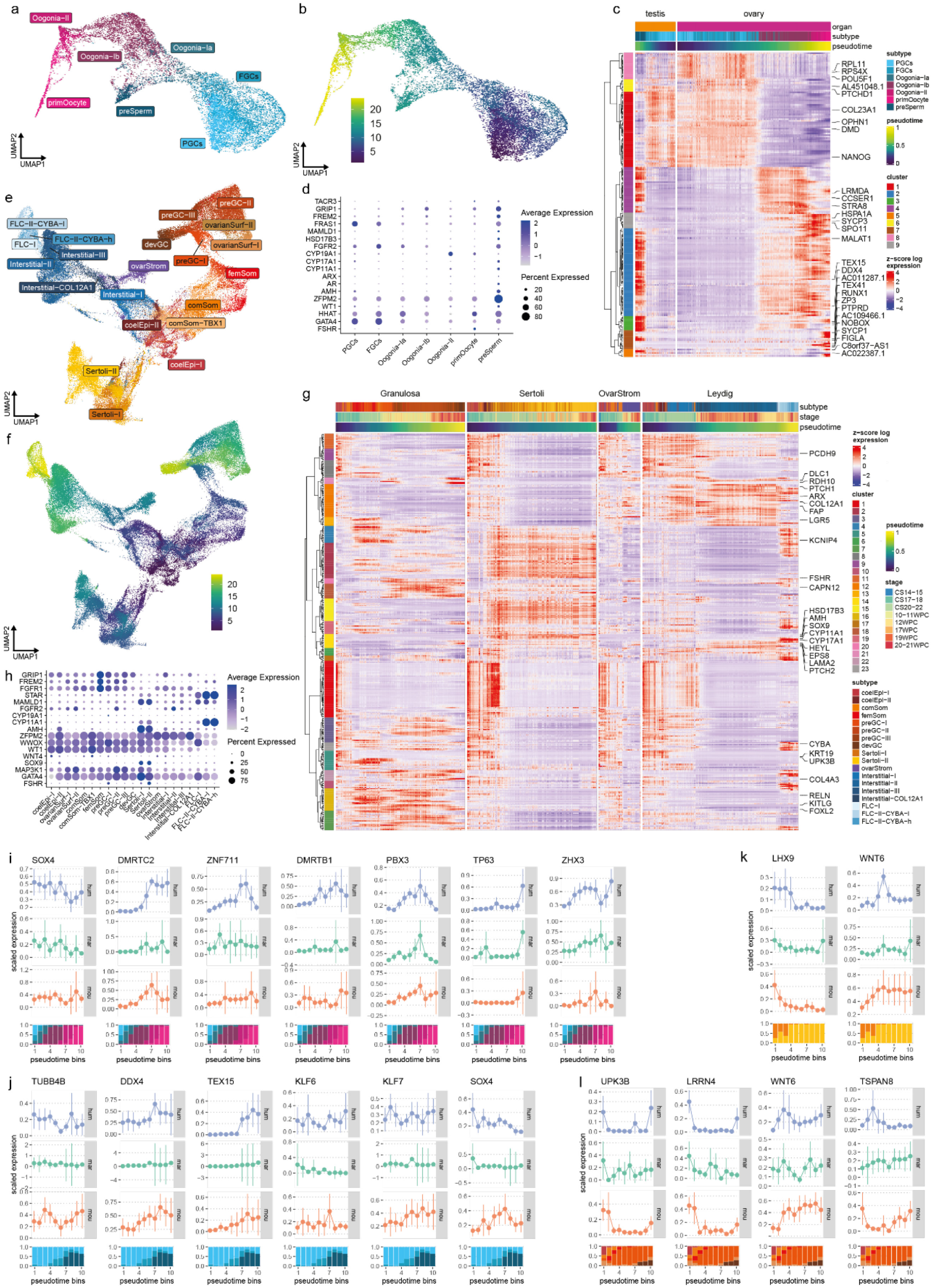
Developmental gene expression trajectories. **a, b,** UMAP embedding of human germ cells coloured by cell state (**a**) or pseudotime value (**b**) based on snRNA-seq data. **c,** Heatmap of gene expression along male and female germ cell pseudotime trajectories. **d**, Dot plot of expression of DSD genes associated with the germ cell trajectory. **e**,**f**, UMAP embedding of human somatic cells coloured by cell type (**e**) or pseudotime value (**f**) based on snRNA-seq data. **g,** Heatmap of gene expression along somatic pseudotime trajectories. **h**, Dot plot of expression of DSD genes associated with somatic trajectories. **i**,**j**,**k**,**l**, Comparisons of expression trajectories of selected genes in female germ cells (**i**), male germ cells (**j**), male supporting cells (**k**), and female supporting cells (**l**) across human, marmoset and mouse.

**Extended Data Fig. 9:**
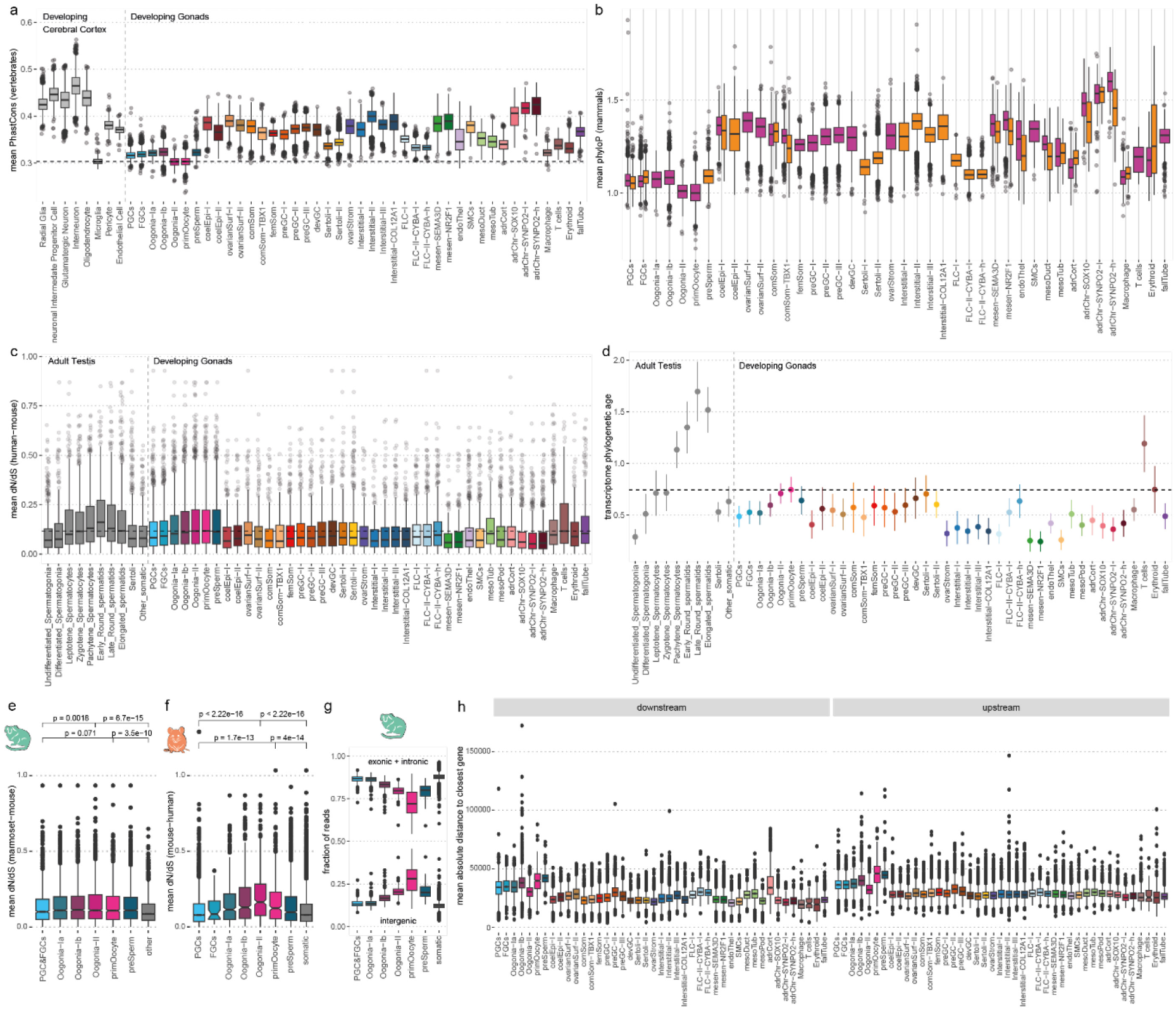
Gonadal cell type evolution. **a**, Sequence conservation of accessible distal CREs in developing human cerebral cortex cell types compared to developing gonad cell types. Published brain data from Trevino et al.^83^ **b**, Sequence conservation of accessible distal CREs in developing gonad cell types separated by sex. **c**, dN/dS based on human-mouse alignment of cell type-specific genes expressed in adult testis compared to the developing gonad. **d**, Phylogenetic age of cell type-specific genes expressed in adult human testis compared to the developing human gonad. **e**,**f**, dN/dS of cell type-specific genes expressed in the developing gonad of marmoset (**e**), based on marmoset-mouse alignment and mouse (**f**), based on mouse-human alignment. Benjamini-Hochberg adjusted *P* values calculated by Wilcoxon rank sum tests. **g**, Fraction of genic (top) and intergenic (bottom) snRNA-seq reads in marmoset germ cell states and all other gonadal cells. **h**, Distance of intergenic snRNA-seq reads to closest gene separated by upstream or downstream genes in the human developing gonad.

## List of Asupplementary tables

**Supplementary table 1:** Overview of snRNA-seq and snATAC-seq libraries.

**Supplementary table 2:** List of marker genes used to annotate cell types in the datasets.

**Supplementary table 3:** DSD genes linked to developmental trajectories in male and female supporting and germ cells.

**Supplementary table 4:** Classification of gene expression trajectories in female germ cells across species.

**Supplementary table 5:** Classification of gene expression trajectories in male germ cells across species.

**Supplementary table 6:** Classification of gene expression trajectories in granulosa cells across species.

**Supplementary table 7:** Classification of gene expression trajectories in Sertoli cells across species.

**Supplementary table 8:** List of species used for dating the emergence of CREs.

